# Evolutionary pathways for the generation of new self-incompatibility haplotypes in a non-self recognition system

**DOI:** 10.1101/260414

**Authors:** Katarína Bod’bvá, Tadeas Priklopil, David L. Field, Nicholas H. Barton, Melinda Pickup

**Affiliations:** Institute of Science and Technology Austria, Am Campus 1, Klosterneuburg A-3400, Austria; Department of Mathematical Analysis and Numerical Mathematics, Faculty of Mathematics, Physics and Informatics, Comenius University, Mlynská Dolina, 84248, Bratislava, Slovakia; Department of Ecology and Evolution, University of Lausanne, UNIL Sorge, Le Biophore, CH-1015, Lausanne, Switzerland; Department of Botany and Biodiversity Research, University of Vienna, Faculty of Life Sciences, Rennweg 14, A-1030 Vienna, Austria

## Abstract

Self-incompatibility (SI) is a genetically based recognition system that functions to prevent self-fertilization and mating among related plants. An enduring puzzle in SI is how the high diversity observed in nature arises and is maintained. Based on the underlying recognition mechanism, SI can be classified into two main groups: self- and non-self recognition. Most work has focused on diversification within self-recognition systems despite expected differences between the two groups in the evolutionary pathways and outcomes of diversification. Here, we use a deterministic population genetic model and stochastic simulations to investigate how novel S-haplotypes evolve in a gametophytic non-self recognition (SRNase/S Locus F-box (SLF)) SI system. For this model the pathways for diversification involve either the maintenance or breakdown of SI and can vary in the order of mutations of the female (SRNase) and male (SLF) components. We show analytically that diversification can occur with high inbreeding depression and self-pollination, but this varies with evolutionary pathway and level of completeness (which determines the number of potential mating partners in the population), and in general is more likely for lower haplotype number. The conditions for diversification are broader in stochastic simulations of finite population size. However, the number of haplotypes observed under high inbreeding and moderate to high self-pollination is less than that commonly observed in nature. Diversification was observed through pathways that maintain SI as well as through self-compatible intermediates. Yet the lifespan of diversified haplotypes was sensitive to their level of completeness. By examining diversification in a non-self recognition SI system, this model extends our understanding of the evolution and maintenance of haplotype diversity observed in a recognition system common in flowering plants.

## 1 Introduction

The origin and maintenance of the extraordinary diversity observed at loci involved in genetically based recognition systems such as MHC in animals (Hedrick 1998), mating types in fungi (May *et al*. 1999) and self-incompatibility (SI) in plants (Wright 1939; Lewis 1949) has long fascinated evolutionary biologists. In all these systems, balancing selection maintains genetic variation (Charlesworth 2006a; Delph and Kelly 2014). Plant self-incompatibility is widespread in flowering plants (Igic *et al*. 2008) and functions to prevent self-fertilization and the consequent deleterious effects of inbreeding depression. Here, negative frequency dependent selection, a form of balancing selection where a rare allele has a selective advantage (Wright 1939), maintains the high diversity observed in nature (Lawrence 2000). Yet one of the most intriguing questions in the evolution of self-incompatibility is how new alleles (S haplotypes) evolve (Uyenoyama *et al*. 2001; Chookajorn *et al*. 2004; Gervais *et al*. 2011). This evolutionary puzzle originates from coevolution of the male and female determining components of the incompatibility reaction. More broadly, many genetic systems involve tightly linked functional components that are kept polymorphic by balancing selection, and that must evolve together. Examples include segregation distorters such as the t-locus in mice (Bennett 1975), supergenes in mimetic butterflies or polymorphic snails, or sex chromosomes (Schwander *et al*. 2014). However, SI systems are an extreme case, in that selection maintains large numbers of alleles, which must be regenerated often enough to avoid loss of variation by random drift. A unifying feature of all SI systems (see the list of acronyms in Table 1) is that the incompatibility reaction is controlled by a single polymorphic locus with low recombination between male- and female-specificities (Takayama and Isogai 2005). A mutation in one component may cause the breakdown of self-incompatibility, with selection against the self-compatible individual due to inbreeding depression (Charlesworth 2006b). This raises the question of whether self-compatibility is involved in the process of S haplotype diversification and if the evolutionary pathways involved vary between different self-incompatibility systems.

An additional complexity in understanding the evolution of SI is that very different evolutionary dynamics are expected for self-recognition (SR) vs. non-self recognition (NSR) self-incompatibility (Fujii *et al*. 2016). Molecular characterization of the three main systems representing the Brassicaceae (pollen ligand and stigmatic receptor kinase), Papaveraceae (Ca2+ dependent signaling) and Solanaceae type, including the Plantaginaceae and Rosaceae (Ribonuclease (SRNase) and F-Box (SLF) protein) (for reviews see Takayama and Isogai (2005), Iwano and Takayama (2012)) has revealed striking differences between self-and non-self recognition systems, especially in the evolutionary relationships between the male- and female-determining components (Fujii *et al*. 2016). In the self-recognition systems (Brassicaceae and Papveraceae), it is the recognition of self that prevents fertilization (Takayama and Isogai 2005; Iwano and Takayama 2012). This results in tight coevolution between male- and female-determining components in a single haplotype shown by their congruent genealogies and a common evolutionary history. However, for nonself recognition (Solanaceae type), the inability to recognize self prevents self-fertilization but allows fertilization by pollen from genetically distinct individuals (Takayama and Isogai 2005; Kubo *et al*. 2010). Here, the male-determining components (SLF) have coevolved to recognize and detoxify the female-determining SRNases from all haplotypes other than their own (Figure 1). High polymorphism and sequence divergence among SRNase genes, compared to high sequence conservation of SLF genes from different haplotypes reflects the different patterns of coevolution in NSR systems (Fujii *et al*. 2016). These inherent differences between SR and NSR systems could result in unique pathways for diversification and for the evolution of novel S haplotypes.

**Figure 1:**
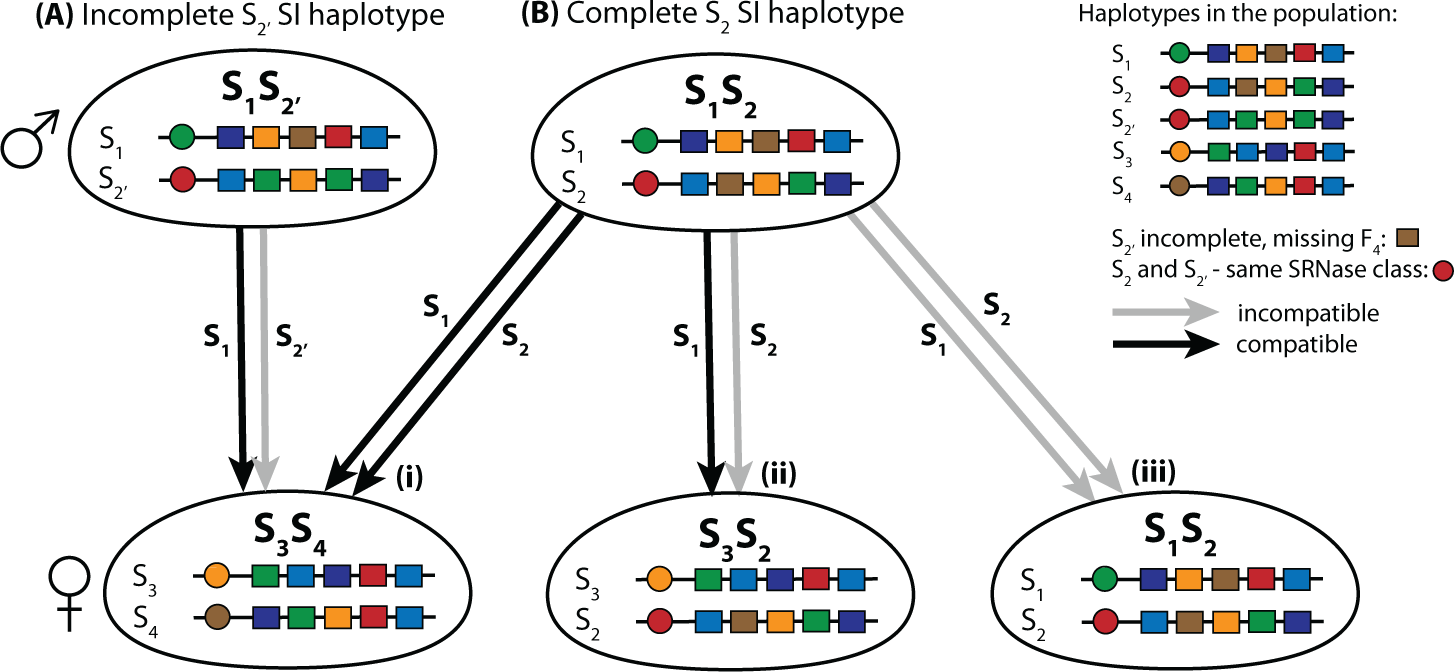
Compatible and incompatible reactions for the gametophytic Solanaceous type non-self recognition self-incompatibility system. The S locus consists of tightly linked female (SRNase) and male (SLF, S locus F-box) determining genes. Circles represent the female SRNase genes (in this example there are altogether *k* = 4 distinct SRNase types) and rectangles the multiple SLF genes. Colours relate to corresponding SRNase and SLF proteins; for example, a green SLF (*F*_1_) is able to detoxify a green SRNase (*R*_1_). A haploid pollen (male) can fertilize a diploid female whenever it has the two SLF that correspond to the two RNase of the female. Each self-incompatible (SI) haplotype is missing the SLF that corresponds to its own SRNase, but has a set of SLF proteins that can recognize and detoxify the SRNases from other haplotypes in the population. A self-compatible haplotype (not shown here) has the SLF corresponding to its own SRNase. Grey lines represent incompatible and black lines compatible reactions. The compatibility reactions in (A) and (B) differ due to a single difference in SLF between *S*_2_, and *S*_2_. Both *S*_2_ and *S*_2_, have the same SRNase and so belong to the same haplotype class. (A) Half compatible reaction due to incompleteness: *S*_2_, is an incomplete haplotype missing one SLF (*F*_4_, brown) for the *R*_4_ SRNase (brown) in the population (completeness deficit 1, see main text for definition). This reduces mate availability so that pollen of type *S*_2_, is not able to fertilize *S*_3_*S*_4_ because it is missing an SLF that corresponds to one of these SRNases. Pollen from *S*_1_ is able to fertilize *S*_3_*S*_4_ (B i) Fully compatible: *S*_2_ and *S*_1_ are both complete haplotypes having the required SLF to detoxify both SRNases in *S*_3_*S*_4_. (B ii) Half compatible: Only *S*_1_ is able to fertilize the female *S*_3_*S*_2_ because a (complete) SI pollen haplotype *S*_2_ is unable to fertilize a female with a haplotype from the same haplotype class. (B iii) Incompatible reaction: No fertilization can occur when both haplotypes of the diploid female and the two male pollen haplotypes are from the same haplotype class - neither pollen haplotype has the SLF required to detoxify either SRNase.

Recent theory has provided a basis for understanding the evolution of novel incompatibility types in both self-recognition and non-self recognition systems. Based on the evolutionary dynamics of self-recognition systems, Uyenoyama *et al*. (2001) and Gervais *et al*. (2011) present a two-step model for the evolution of novel S haplotypes where first there is a mutation in the male specificity, followed by a corresponding mutation in the female specificity. Under conditions of strong inbreeding depression and lower selfing rate, Uyenoyama *et al*. (2001) found that new specificities arise via evolutionary pathways that include a loss of self-incompatibility (i.e. a self compatible intermediate). However, the novel S haplotype was often found to replace the ancestral haplotype, so that diversification an increase in haplotype numbers - was limited. Moreover, in this model, diversification through the pathway that involved an initial female (pistil) mutation followed by a mutation in the male (pollen) component was thought to be impossible (Uyenoyama *et al*. 2001). This model was analyzed further by Gervais *et al*. (2011) who found that diversification was possible under conditions of high inbreeding depression and a moderate to high rate of self-pollination.Furthermore, the rate of diversification was higher with fewer S haplotypes in the population (Gervais *et al*. 2011) since there is stronger selection when there are fewer S haplotypes.

For non-self recognition systems, Fujii *et al*. (2016) recently presented a conceptual two-step model for novel S haplotype evolution. In this model, a mutation to generate a new SLF gene occurs first, which, given there is no associated cost, increases in frequency via drift. Once the novel SLF gene is common enough, a self-incompatible haplotype can be generated by a corresponding mutation in the SRNase in another haplotype. Fujii *et al.* (2016) then provide simulations to support this model that suggest diversification via this pathway is possible, but that genetic exchange is important for new haplotype evolution (Fujii *et al.* 2016). Although this model provides a useful basis for investigating the evolution of novel S haplotypes in NSR systems, there are questions regarding its plausibility as the driving force in generating novel SI haplotypes. Firstly, we observe that for the newly generated SI haplotype to be able to invade, the novel SLF gene created in the first step must increase to a high frequency as otherwise the fertilization of the new SI haplotype will be unlikely. This implies that the new SLF gene should ideally occur in many haplotypes, and therefore either recurrent mutations must happen on different haplotypes, or genetic exchange must be common. Second, even if the new SLF spreads to reach a sufficient frequency, the newly created incomplete haplotype is still rejected by at least two haplotypes, itself and the ancestral type, and is thus at a selective disadvantage due to mate limitation. Finally, their model also only considers the evolution of new incomplete haplotypes through a pathway that maintains SI. Therefore, it remains unclear to what extent, if at all, the path proposed by Fujii *et al.* (2016) is responsible for the diversification of SI haplotypes in NSR systems. Further theory is therefore required to examine all potential evolutionary pathways, including those with self-compatible intermediates, and examine how the evolutionary outcomes vary with parameters such as inbreeding depression, selfing rate, number of haplotypes and with mate availability mediated by incompleteness.

Here we combine analytical theory with simulations to examine the interplay between completeness, drift and mutation for diversification in a NSR SI system. First, we propose an explicit population genetic model for the evolution of novel S haplotypes for non-self recognition systems that includes pathways with a maximum of two selected (non-neutral) mutations (Figure 2). In this model there are five potential evolutionary pathways, all starting from an ancestral SI haplotype which we arbitrarily label *S*_*k*_. This haplotype may be in two states, with either the presence 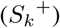 or absence 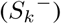 of the novel SLF, so that a single neutral mutation separates the two states. Following this, the first non-neutral mutation may be a change in the SRNase (female specificity) gene (pathway 1 from 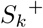 or pathway 5 from 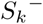), the simultaneous loss and gain of an SLF (male specificity) gene (pathway 2), or gain of an SLF gene (pathway 3 from 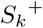 or pathway 4 from 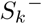) (Figure 2 A,B). The second non-neutral mutation then involves either the gain of the novel SRNase (pathways 2, 3 and 4; notice that pathway 3 requires an additional neutral mutation), loss of the novel SLF to restore SI (pathway 1), or gain of the ancestral SLF (pathway 5). Consequently, in this model, a new S haplotype can evolve via a number of pathways, including four that involve a self-compatible intermediate, and one in which SI is maintained. Genetic and demographic factors such as selfing rate, level of inbreeding depression, mutation rates, number of haplotypes and population size (Wright 1939; Uyenoyama *et al*. 2001; Gervais *et al*. 2011) may affect the rate and trajectory of diversification at the S locus. Despite their potential importance, to date, no models have examined these conditions and how they interact with S haplotype diversification in NSR systems. Our model examines these conditions and relates this to the different evolutionary pathways that may generate novel S haplotypes. Moreover, by considering all pathways, and reducing our model to correspond to previous models (Uyenoyama *et al*. 2001; Gervais *et al*. 2011), we can also directly compare these studies (based on SR) with ours, which considers the dynamics of NSR SI systems.

To investigate the pathways and conditions associated with novel S haplotype evolution, we first establish a deterministic population genetic model for the NSR system. By examining all potential evolutionary pathways (Figure 2), we ask: what conditions (inbreeding and self-pollination rate) are associated with the evolution of new S haplotypes and does this vary with evolutionary pathway? Do novel S haplotypes evolve through self-compatible intermediates? How does completeness (having a full set of SLFs from the population) influence diversification? And, does this vary with haplotype number? Following this, using stochastic simulations with finite population size, we ask, in addition to the effect of inbreeding depression and selfing rate, what is the influence of population size, mutation rate and number of potential haplotypes on the evolution of new specificities in a NSR system? Finally we ask if certain evolutionary pathways are more common for SI haplotypes with long lifespans and high frequencies. We found that diversification was possible in our analytical model, but the parameter space for diversification was more limited than in stochastic simulations and varied with haplotype number and the level of completeness. Furthermore, in the analytical model, diversification generally resulted in a short-term increase in S haplotype numbers, especially in the pathway that maintains SI. In comparison, in the stochastic simulations, S haplotype diversification was observed even for the pathway maintaining SI. This difference highlights the potential importance of drift, unconstrained mutational order and mutation rate for diversification outcomes. By considering all evolutionary pathways, and the conditions associated with S-haplotype diversification, our model combines analytical theory and stochastic simulations to provide new insights into the evolutionary dynamics of non-self recognition self-incompatibility systems in plants.

**Figure 2:**
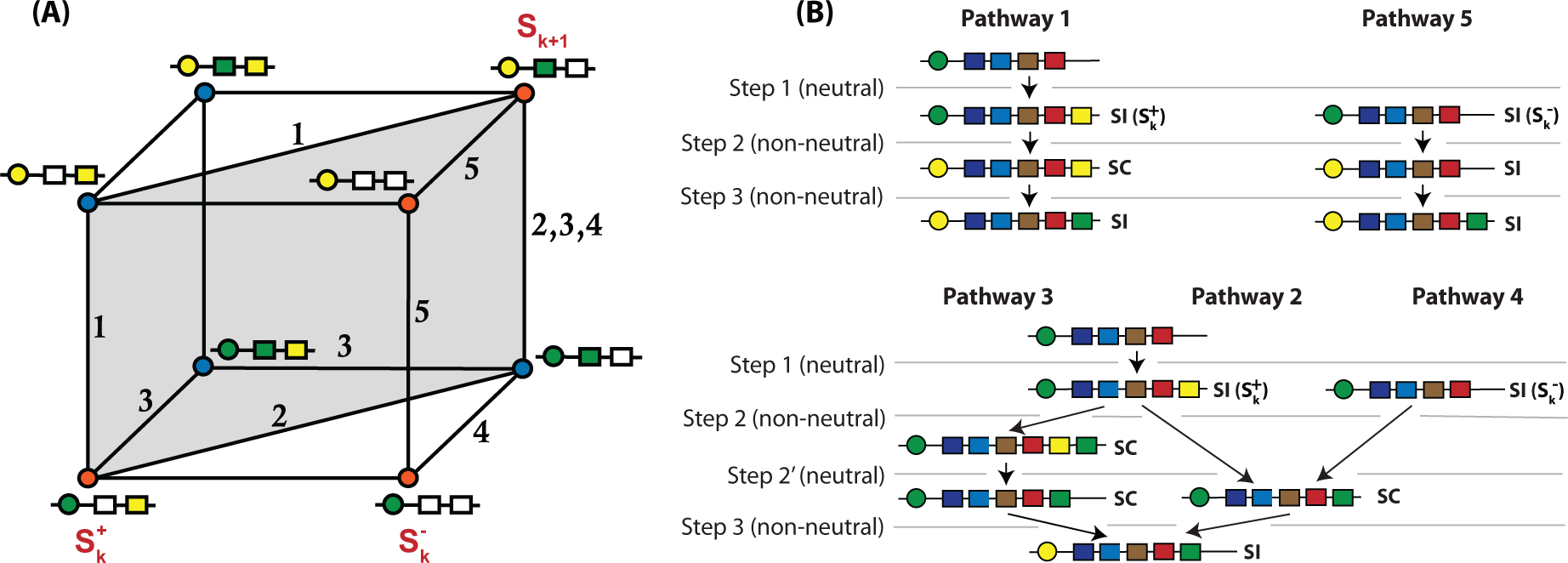
(A) Potential pathways for the evolution of a novel self-incompatible (SI) haplotype for the gametophytic SRNase/F-box non-self recognition self-incompatibility system. A novel SI haplotype in class *S*_*k*+1_ (top right corner) evolves from an ancestral self-incompatible haplotype in class *S*_*k*_ (either 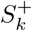 bottom left corner or 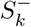 bottom right corner; the transitions between 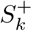 and 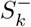 are selectively neutral, see main text) via a number of potential evolutionary pathways. Each side of the cube represents a mutation in either the male (SLF; represented by a box) or female (SRNase, represented by a circle) specificities that make up the S haplotype. Mutations along the horizontal planes of the cube involve the male SLF gene, with a deletion (from a filled to open box) or addition (from open to filled box). An open box may be seen either as an SLF that is not utilized, or as a place-holder for a SLF which is then added for example by duplication (see the main text). Mutations in the vertical plane represent a mutation to generate a novel SRNase (from green to yellow circle). At each vertex, red circles represent self-incompatible haplotypes (no SLF to detoxify its own SRNase) and blue circles self-compatible haplotypes (SLF present to detoxify its own SRNase). 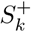 is the initial haplotype with the SLF corresponding to the novel SRNase; while in 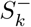 the SLF for the novel SRNase is missing. Pathways 1 and 2 (starting with 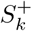, or alternatively with 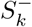 followed by a neutral mutation to obtain 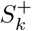) involve two mutations with a self-compatible (SC) intermediate. Pathway 5 (starting with 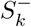) involves two mutations and self-incompatibility (SI) is maintained. Pathway 3 (starting with 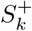) involves three mutations and a SC intermediate. Pathway 4 (starting with 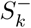) involves two mutations and a SC intermediate. The highlighted grey diagonal rectangle indicates pathways 1 and 2 which have been previously considered by Uyenoyama *et al*. (2001); Gervais *et al*. (2011). (B) Mutations involved in the five pathways that result in a novel SI haplotype. For all pathways, the first mutation (step 1) is neutral because there is not yet a corresponding SRNase in the population. This step involves either the presence 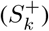 or absence 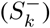 of the novel SLF. Step 2 is a non-neutral mutation that involves either the gain of the novel SRNase (pathways 1 and 5) or gain of the ancestral SLF (pathways 2 and 4). For pathway 3, this occurs in two-steps, with the gain of the ancestral SLF, then the loss of the novel SLF. The final non-neutral mutation (step 3) involves either the loss of the novel SLF (pathway 1), gain of the ancestral SLF (pathway 5) or gain of the novel SRNase (pathways 2, 3 and 4).

## 2 Methods

### 2.1 The non-self recognition system

The genetic mechanism of our NSR system builds on the collaborative non-self recognition model proposed by Kubo *et al*. (2010); Fujii *et al*. (2016). This is based on a gametophytic SI system where incompatibility is determined by the haploid genotype of the pollen. We assume that each haplotype consists of two tightly linked loci, an R-locus and an F-locus. An R-locus determines the female specificity and consists of a single RNase *R*_*i*_, whereas at the F-locus male specificity is determined collaboratively by several SLF genes (F-boxes) *F*_1_, *F*_2_,…. In the analytical model (see below) we don’t make any assumptions on the length (*L*) of this sequence, and so the appearance of a new SLF may either keep the length of the haplotype *L* intact (e.g. a change of specificity of an old SLF), or *L* may increase due to a duplication event. However, in the stochastic simulations (see below), we assume that the number of SLF in each haplotype is constant (*L* is constant), implying that a mutation in the SLF results in a simultaneous appearance (addition) and disappearance (deletion) of an SLF. We assume a one-to-one correspondence between SLFs and RNase, such that each SLF *F*_*i*_ is assumed to detoxify (recognize) exactly one RNase *R*_*i*_ and each RNase R_i_ is recognized by exactly one SLF *F*_*i*_ This implies, for example, that a pollen haplotype needs to have both SLFs *F*_*i*_ and *F*_*j*_ in order to recognize, and thus be able to fertilize, a diploid plant with *R*_*i*_ and *R*_*j*_ RNase (Figure 1). Conversely, incompatibility results from the failure of a pollen grain to recognize both RNase of a diploid female. For a pollen grain to be able to fertilize all individuals in the population it needs to have as many SLFs as there are RNase present in the whole population. If a haplotype has all the required SLFs we call it *complete* (and otherwise *incomplete)*. We emphasise that completeness is a property of a haplotype, but also depends on the composition of the population (Figure 1). The level of incompleteness of a haplotype is described by the completeness deficit which measures the number of missing SLFs. A *haplotype class* represents all haplotypes that have the same female specificity (RNase) but may have different sets of SLFs. We treat as one SI class all complete and incomplete SI haplotypes with the same RNase (Figure 1). and so the appearance of a new SLF may or may not be accompanied by the increase of the number of SLF. For example

**Table 1:** Abbreviations and terminology.

| Term | Explanation |
| --- | --- |
| SI | Self incompatible/Self incompatibility |
| SC | Self compatible/Self compatibility |
| NSR | Non-self recognition |
| SR | Self recognition |
| SLF, F-box | S-Locus, F-box |
| R-locus, F-locus | Female/male part of the S-locus |
| SRNase | S-Locus (haplotype specific) RNase |
| Complete haplotype | Haplotype that can fertilize every SI individual in the population |
| Completeness deficit | The number of non-self SI haplotype classes that a pollen grain cannot fertilize |
| Haplotype class | All haplotypes with the same female specificity (RNase) |
| NFDS | Negative frequency dependent selection |

### 2.2 Evolutionary pathways for the generation of new SI haplotypes: The Cube

Due to the collaborative nature of the NSR system, the generation of a new SI haplotype not only requires mutational changes on a focal haplotype but also in other haplotypes in the population. Moreover, some of the required mutations may be under functional selection, but some are not. We thus need to make a distinction between the number of neutral and non-neutral (selected) mutations in the focal haplotype, as well as the numbers on other haplotypes in the population. In this paper we consider all evolutionary pathways for the generation of new SI haplotypes that allow up to two selected mutations on a single haplotype. As in the previous work on the evolution of new SI haplotypes (Uyenoyama *et al*. 2001; Gervais *et al*. 2011), we start by considering an initial population of k complete SI haplotype classes, and discuss all the mutations that lead to a new complete SI haplotype class.

#### Step 1 (initial condition; neutral mutations)

For every diversification pathway the first mutation in the population must necessarily give rise to a haplotype with a novel male specificity, i.e. a new SLF; a haplotype with a new female specificity will never be fertilized and thus can never invade (Figure 2). We start our analysis by either assuming that the novel SLF already exists in some SI haplotype classes in the population, or that it fixes due to drift. And so we enter the Cube (Figure 2).

#### Step 2 (1st non-neutral mutation)

Suppose that the novel not-yet utilized SLF, which we label *F*_*k*+1_, is fixed within n different SI classes. We will follow changes on a haplotype, say *S*_*k*_ (or any other of the *k* existing SI classes; note that *S*_*k*_ is not necessarily the class that underwent the previous diversification event), which may or may not have the novel SLF *F*_*k*+1_. We write 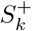 if the haplotype class has the novel SLF *F*_*k*+1_, and 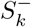 if not. Each pathway is described in Figure 2. Three selectively different events may follow: a mutation in the R-locus, either (i) on a 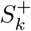 haplotype (path 1), or (ii) on a 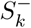 haplotype (path 5), or a mutation in the F-locus (iii) such that haplotype *S*_*k*_ (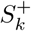 or 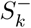) obtains an SLF *F*_*k*_ that corresponds to its RNase *R*_*k*_ (paths 2, 3 and 4).

#### Step 3 (2nd non-neutral mutation)

The final step requires either gain of an SRNase (pathways 2,3,4 - notice that path 3 requires an additional neutral mutation), gain of the ancestral SLF (pathway 5), or replacement of the new SLF by the ancestral SLF (pathway 1); see Figure 2.

#### Final state of the population

All five evolutionary paths lead to a novel complete SI haplotype *S*_*k*+1_. If all initial *k* SI haplotype classes remain in the population during this process, we say that the paths are diversification pathways. Recall, however, that since we assume that in *k* - *n* of the initial *k* SI classes *F*_*k*+1_ is absent, then after the diversification process (provided no further mutations have happened in the population), *k* - *n* SI classes are incomplete and will not be able to fertilize the new SI class *S*_*k*+1_.

Two remarks are in order. Firstly, the steps described above give a detailed description of all the changes in the population needed to generate a new complete SI class. This, however, may seem an overly strict requirement, especially for large *k*, since missing one or two SLFs affects only slightly the frequencies of haplotypes compatible with the focal haplotype. If we relax the requirement of completeness, we only need, in principle, two changes (one non-neutral) in the population. First, a single haplotype class gains a ”not-yet utilized” SLF *F*_*k*+1_, after which another haplotype class *S*_*k-*_ undergoes a mutation in the R-locus generating an incomplete SI haplotype *S*_*k*+1_ (step 2 1st non-neutral mutation of path 5, Figure 2). This path was also identified by Fujii *et al*. (2016). It remains to be seen, however, whether this path is likely (or even possible) under the various levels of incompleteness (parameter n) and demographic parameters used in this paper. Secondly, if we assume that all *k* SI classes initially have the new SLF *F*_*k*+1_ (i.e. *n* = *k*), and then restrict the subsequent mutations to the diagonal of the cube (highlighted rectangle in Figure 2, i.e. considering only paths 1 and 2), we recover the SR model in Uyenoyama *et al*. (2001) and Gervais *et al*. (2011). This allows us to directly compare the evolutionary diversification pathways in NSR and SR systems. Finally, we remark that since in the NSR model pathways 1 and 2 imply a simultaneous loss and gain of two specific SLF, we predict that these pathways (if feasible) are less common than the alternative pathways 3,4 and 5. This should be observed in the stochastic simulations where the rate of mutations is considered explicitly. In the next section, we present the life cycle of an individual and give the population genetic equations for infinite and finite population sizes. We then detail the procedures and definitions used in stochastic simulations.

### 2.3 The life cycle of an individual and the dynamics of the population

In this section we first give the life cycle of an individual and then give the recurrence equations to study the dynamics of haplotypes for both infinite and finite population models. The infinite population model is the large population limit of the stochastic finite population model.

Consider a well mixed population with non-overlapping generations, such that one iteration of the (finite and infinite population) model represents the full life-cycle of an individual (e.g. annual plants). At the beginning of the season, each diploid individual plant produces, and receives, a large number of haploid pollen grains. Of all the pollen received a proportion α is assumed to come from the same individual and 1 - α from a pollen pool from all the other plants (i.e., global pollen dispersal). The proportion of haplotypes received is proportional to the frequency distribution of the pollen in the whole population. Importantly, we assume that all outcrossed mating events are among unrelated individuals and that self-fertilized offspring undergo inbreeding depression Uyenoyama *et al*. (2001); Gervais *et al*. (2011). Self-fertilized offspring survives until adulthood with a fixed probability 1 - δ relative to outcrossed offspring. After offspring are produced all adults die. A possible mutation occurs at the time of reproduction. We assume no recombination in the S-locus. In this paper we consider no geographic structure. Thus we make no distinction between globally and locally incomplete S haplotypes, implying that mate limitation due to completeness plays a lesser role in structured populations (see Discussion).

#### Infinite population model

Following the assumptions of the life cycle of an individual (see above), the probability that an individual with genotype *g* is self-fertilized is equal to the number of self-pollen grains received that are self-compatible, divided by the total number of compatible (self and non-self) pollen grain received:

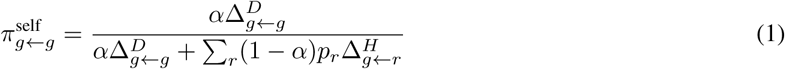

where 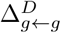 is a diploid fertilization function that gives the fraction 0, 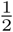 or 1 of self-pollen that can self-fertilize, 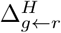 is a haploid fertilization function that gives the probability that a pollen with haplotype *r* can fertilize a diploid individual *g*, and *p*_*r*_ is the frequency of (haploid) pollen *r* in the whole population. Similarly, the probability that an individual with genotype *g* is outcrossed with haploid pollen *h* is

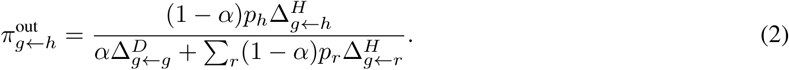

If the plant is self-fertilized, the offspring survives until adulthood with probability 1 - δ relative to outcrossed offspring. The frequency of (diploid) genotype *r* in the next generation is

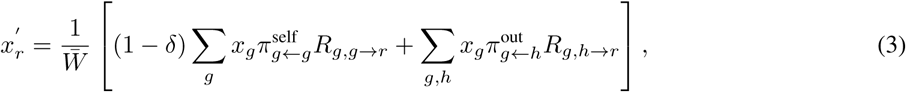

where 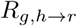 is the probability that a diploid individual g that is fertilized by a haploid pollen h produces a diploid offspring *r* (according to the Mendelian rules), 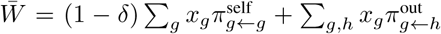 is the average fitness in the population, and where for clarity we use *x* to denote (diploid) genotype frequencies.

To study the conditions for the various diversification pathways we suppose, in the infinite population model, that initially the population consists of *k* SI classes, with equal frequencies 1/*k* and no SC haplotypes. Moreover, *n* out of these *k* classes have the novel not-yet utilized SLF *F*_*k*+1_ fixed within the class. The haplotypes in the remaining *k* - *n* classes do not have *F*_*k*+1_. We note that each haplotype may have a different number of SLF, i.e. *L* is not fixed in the infinite population model.

#### Finite population model

Let *N* denote the number of individuals considered in each simulation (i.e. 2*N* haplotypes). Each haplotype is represented by a sequence

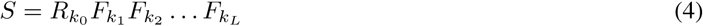

denoting the states at a single female determining SRNase and a fixed number *L* of male SLF genes. Keeping *L* fixed implies that the appearance of a new SLF gene is accompanied by loss of another SLF gene. However, the effect on fitness becomes small when *L* is large as it increases the likelihood that the lost SLF gene doesn’t correspond to any of the RNase in the population and thus has no affect on the mate-availability of the haplotype. Unlike in the infinite population model, where we assumed initial presence of SI classes, the initial state in the finite population simulations consists of identical self-compatible haplotypes, i.e., all haplotypes carried the same SRNase and the same (random) sequence of SLF-genes. This choice of the initial state allowed us to capture the initial phase of the evolution of SI haplotypes where self-incompatibility appears and forms a functional system with at least three SI haplotypes. Moreover, the combination of mutation and drift results in intermediate populations which can be much more diverse in comparison with the infinite population model, i.e., the finite population model allows multiple simultaneous diversification events at the same time. Thus the finite population model may reveal some of the limitations of the infinite population model. Each generation consisted of the life-cycle described above; while each life cycle consists of mutation, followed by reproduction and viability selection.

##### 1. Mutation

We assumed a finite space of distinct RNase types {*R*_1_, *R*_2_,…, *R*_*nR*_} and SLFs {*F*_1_, *F*_2_,…, *F*_*nF*_} where the pollen type *F*_¿_ targets the RNase *R*_*i*_ (*n*_*F*_ = *n*_*R*_). The number of mutations in the RNase and in the SLFs were randomly drawn from a Binomial distribution with parameters (2*N, μ*_*R*_) for RNase mutations and (2*NL, μ*_*F*_) for SLF mutations and randomly placed on the genotypes. Here *μ*_*R*_ and *μ*_*F*_ represent the probability of a mutation in a given generation either in the RNase or in a single SLF. The Binomial distribution can be replaced by the Poisson distribution with parameters 2*Ν μ*_*R*_ (female) and 2*NL μ*_*F*_ (male), since the two distributions are the same in the limit of small *μ*_*R*_, *μ*_*F*_ and large *N, L* (valid throughout our simulations). Mutation rate was the same for all RNase-mutations *R*_*i*_ → *R*_*j*_, and similarly for all SLF mutations *F*_*i*_ → *F*_*j*_ for any *i, j*.

##### 2. Reproduction and viability selection

The second part of the simulation consisted of randomly generating the mating partners and their offspring, incorporating the compatibility between individuals and selection in terms of selfing α and inbreeding depression δ.

In the infinite population model the frequencies of individual genotypes *g* in the next generation can be obtained by solving a system of difference equations (3). However, stochastic simulations use probabilistic rules to determine offspring production for all parent combinations. These rules quantify the probability that a female with a genotype *g* and a male with a genotype *h* produce an (adult) offspring as

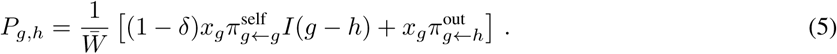

where 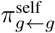 and 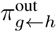 are defined in (1)-(2) and *I* (*g* - *h*) = 1 when *g* = *h* and 0 otherwise. The mean population fitness 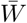 ensures that 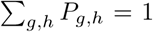. The first term in (5) reflects outcrossing events between genotypes *g* and *h*, while the second term in (5) captures self-fertilization and is nonzero only when *g* = *h*. The simulation required two steps:

- Using the current genotypes {*g*_*i*_}, their frequencies *x*_*gi*_ and compatibility relationships 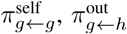 we first generated N parental pairs *g, h* (distinguishing between females g and males h) by stochastic sampling with probabilities (*g, h*) → *P*_*g,h*_.
- Next, we randomly generated a single offspring for each parental pair using Mendelian inheritance.

We ran the simulation for a fixed number of generations (10^3^ - 10^4^ generations) with parameters *N, μ*_*R*_, *μ*_*F*_, α, δ, summarized in Table 2. The structure of the space of possible haplotypes depends on our choice of parameters *n*_*R*_, *n*_*F*_ and *L*. We recorded the list of genotypes in the population at all times, as well as a list of all mutations. This allowed us to trace the key measures and to:

1. Classify haplotypes based on their haplotype classes. Initially, a single SC class is present in the population (no SI classes were initially present).
2. Classify haplotypes based on compatibility among SI classes. We distinguish between SI and SC haplotypes and, in combination with the grouping in 1, we plot the frequencies of SI and SC haplotypes within each class.
3. classify haplotypes based on completeness defict,i.e the number on non-self SI classes that cannot be fertilized because of missing SLFs. This measure depends on the number of SI classes,as it cannot exceed the number of SI classes-1. The minimal completeness defict of an event is computed as a minimum deficit through all times during the life time of the event and through all SI haplotypes within the class at each time.
4. Identify all birth/death events of the SI classes. Birth of a new SI class of type *k* is an event which occurs at generation *t* if there is at least one SI haplotype with SRNase *R*_*k*_ at the *t*-th generation while there was none such haplotype in the previous generation. Death of the *k*-th SI class occurs when the last SI haplotype from that SI class vanishes at generation *t*, provided there was at least one such individual at generation *t* - 1.
5. Trace the genealogy of the SI classes. We can trace the order of mutations that led from the ancestral haplotype to any other haplotype and record the times at which the mutation occured. This allows us to trace the pathway leading to a birth (or death) event of any SI class. We chose the representative haplotype of each event as the first haplotype that reached the minimal completeness deficit of this event. We found that the results were almost identical when the last haplotype in the class was chosen as a representative haplotype. We then traced back all its ancestors from the current and previous RNase class and projected it onto a mutation cube in Figure 2.

**Table 2:** Parameters of the NSR model used in simulations.

| Parameter | Description | Range of values |
| --- | --- | --- |
| $N$ | Population size | $\{200, 1000\}$ |
| $\alpha$ | Self-pollination rate | $[0, 1]$ |
| $\delta$ | Inbreeding depression | $[0, 1]$ |
| $\mu_R$ | Mutation rate of SRNase | $\{10^{-3}, 10^{-4}\}$ |
| $\mu_F$ | Mutation rate of SLFs | $\{10^{-3}, 10^{-4}\}$ |
| $L$ | Number of SLFs in a haplotype | 15 |
| $n_R$ | Number of possible SRNase | $\{15, 50\}$ |
| $n_F$ | Number of possible SLFs | $\{15, 50\}$ |

## 3 Results

### 3.1 Theoretical predictions of evolutionary outcomes

Our model examines five potential pathways for the evolution of new SI haplotypes (see Figure 2), which includes four pathways with self-compatible intermediates (pathways 1, 2, 3 and 4) and one where self-incompatibility is maintained (pathway 5, see Figure 2). These pathways are also defined by their starting condition (i.e. before the first functional mutation occurs), with pathways 1, 2 and 3 starting with the novel SLF 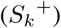, while the novel SLF is initially absent in pathways 4 and 5 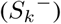. The transition between the two starting positions is selectively neutral and requires only a single mutation in the SLF. We divide the parameter space, proportion of self-pollination (α) and inbreeding depression (δ), into regions that represent; diversification (green; long-term increase in the number of SI haplotypes, loss of the intermediate mutant), short-term diversification (light magenta; short-term increase in the number of SI haplotypes), exclusion of the novel SI haplotype (*S*_*k*+1_; grey), novel SI haplotype *S*_*k*+1_ replaces its ancestral SI haplotype (red; no diversification), SC haplotypes go to fixation (below the thick line; no SI present in the system), see Figure 3.

#### The maintenance of complete and incomplete haplotypes

To disentangle the complexities involved in the diversification process, it is first useful to compute the effect of (in)completeness (number of missing SLFs) on mate-availability and fitness. Complete SI haplotypes are maintained in the population due to NFDS, and when rare, they have a selective advantage and increase in frequency. However, this is not necessarily true in a population with incomplete haplotypes. Here, rare incomplete haplotypes are expected to have a fitness advantage over common incomplete haplotypes with an equal, or lower, level of completeness, but not over haplotypes with higher level of completeness (e.g. fully complete haplotypes). Therefore, even though incomplete haplotypes are under NFDS, they may not have sufficiently high fitness to increase in frequency when rare. We then aim to resolve the conditions under which haplotypes with varying levels of completeness are maintained in the population. This is useful for investigating the feasibility of diversification pathways because, in general, diversification will result in some fraction of incomplete haplotypes.

Aligned with our assumptions on the diversification pathways (see Methods), we provide the exact analytical calculations for which *k* + 1 SI haplotypes, some complete and some incomplete, can coexist in the population. Out of the *k* + 1 haplotypes *k* - *n* are assumed incomplete and lack a single but identical SLF. Following this, *n* +1 are assumed complete SI haplotypes, including one which can’t be fertilized by the incomplete haplotypes, which corresponds to the haplotype *S*_*k*+1_ in the diversification pathways (Appendix A). We found that when there were, in addition to *S*_*k*+1_, no complete haplotypes (*n* = 0) coexistence is only possible when the initial number of haplotypes (k) is three, but not for *k* > 3. With one complete haplotype (n = 1) coexistence is only possible for 3 ≤ *k* ≤ 6 but never for *k* > 6. However, when all haplotypes are complete (*n* = *k*), coexistence is possible for all k. This implies that diversification is only possible when either a single haplotype is complete (*n* = 1) but the total number of haplotypes is small (*k* ≤ 6) or when all haplotypes are complete (*n* = *k*). Despite these rather strict requirements for diversification there are several caveats. Firstly, we found that even if the incomplete haplotypes are not maintained in the population and eventually go extinct, an incomplete haplotype that is initially common will decreases to zero frequency only very slowly and could thus be rescued from extinction by gaining the missing SLF. Secondly, these results do not explicitly consider diversification events that involve the coexistence of SI and SC types. Thirdly, we only considered the case where all incomplete haplotypes lack a single and identical SLF, thus coexistence may be possible for haplotypes with varying levels of completeness and different missing SLFs. Finally, our results hold only for global pollen dispersal. When dispersal is local, we expect that globally incomplete haplotypes that are locally complete will be maintained in the population. Next, we complement these results by studying all evolutionary diversification pathways, including intermediate haplotypes, to determine which diversification pathways are the most likely.

#### Pathway 1: SRNase as the first mutation

Previous models (Uyenoyama *et al*. 2001; Gervais *et al*. 2011) suggested that diversification is not possible through pathway 1 (first mutation in the female-specificity). Our model can be reduced to correspond to these models for pathways 1 and 2 when all SI haplotypes are assumed complete and the dynamics are constrained to the diagonal of the cube (highlighted pathways in Figure 2). Pathway 1 represents a mutation first in the female SRNase leading to a SC intermediate followed by the pollen-part mutation (SLF) to produce a SI haplotype (Figure 2). Our results show that diversification via this pathway is possible for all levels of completeness (*n*) and haplotype number (*k*) (see Figure S3 in the Supplemental Information). This apparent discrepancy with our previous result that diversification is possible only for *n* = 1 or *n* = *k* originates from the fact that here, in pathway 1, the intermediate SC haplotype is not excluded in the diversification process. However, when considering diversification through all pathways (Figure 2), diversification via pathway 1 occurs in the parameter region where SC haplotypes (produced in pathways 2, 3 and 4) have a fitness advantage, ultimately resulting in the loss of SI (region below the thick black line in Figure 3, see also Figure S3 in the Supplemental Information). Consequently, for both SR and NSR systems, it appears that diversification is unlikely through this evolutionary pathway.

#### Pathways 2 to 5: the effect of completeness and haplotype number

We found that for pathways 2 to 5 completeness interacts with haplotype number to determine the values of self-pollination and inbreeding depression where diversification is possible. When all haplotypes were incomplete (*n* = 0) diversification was not possible through any of the pathways, independent of the number of haplotypes in the population (Figure 3i). In contrast, for example, when *k* = 5, long-term diversification occurred under conditions of high self-pollination rates and inbreeding depression under two scenarios; firstly, through pathways 2, 3 and 4 when the number of complete haplotypes was one (n =1; green region in Figure 3ii) and secondly, when all haplotypes were complete, through pathways 2 and 3 (green region in Figure 3iv). When the number of complete haplotypes was between 1 and *k*, only short-term diversification was possible (light magenta region in Figure 3iii), and this occurred though pathways 2, 3, 4 (SC intermediates) as well as pathway 5 (SI maintained). Because all intermediate SC haplotypes are excluded after diversification, in the short-term, new SI haplotypes coexist in the system following diversification, but all incomplete haplotype classes then slowly go extinct (see the results above). In this case, extinction is only prevented by mutations that result in incomplete haplotype classes becoming complete (obtaining all SLF genes in the population).

The effect of completeness on diversification was also observed at higher haplotype numbers. When *k* = 8, there was no diversification when n =1 (the condition for possible diversification when *n* =1 is *k* < 7, Appendix A), with the novel SI haplotype (*S*_*k*+1_) unable to invade the population (Supplemental information Figure S6). This is in contrast to diversification in the narrow region of self-pollination and inbreeding depression when there are fewer haplotypes (*k* = 5; Figure 3Aii vs. Supplemental Information Figure S6ii). Yet, similar to when *k* = 5, short-term diversification was observed when the number of complete haplotypes were between 1 and *k* (Figure 3iii and Supplemental Information Figure S6iii), and long-term diversification at high self-pollination (α) and inbreeding depression (δ) when all haplotypes were complete (Figure 3iv and Supplemental Information Figure S6iv). This suggests that the conditions for diversification are restricted to a narrow region of α and δ, but that this is more flexible when there are fewer haplotypes (smaller k), as diversification can occur at both low and high levels of completeness. Moreover, completeness will determine if there is long- or short-term coexistence of novel haplotypes in the population. The parameter spaces for diversification for each evolutionary pathway across a broader range of haplotype numbers and levels of completeness are outlined in Appendix A-C.

If the selfing rate α and inbreeding depression δ are constant, the SR model of Uyenoyama *et al*. (2001) and Gervais *et al*. (2011) predicts very low numbers of S-alleles. This is because the diversification regions from *k* to *k* +1 S-alleles generally do not overlap, unless *k* is small and thus in most cases only a single diversification event is possible. Interestingly, the diversification regions for the infinite population NSR model are identical to those for the SR model, see Figure 2 in Gervais *et al*. (2011). The similarity between the SR and NSR models is because all haplotypes are initially assumed complete (in both SR and NSR models) before the novel female specificity arrives in the population. We show the diversification regions in Figure 3 for *k* = 3 - 8 S-alleles. Multiple diversifications may occur only when the diversification regions overlap (grey color in Figure 3B). For example, grey region I allows diversification from *k* = 3 to *k* = 5 since it is the intersection of the regions for *k* = 3 and *k* = 4. Similarly, region II allows diversification from *k* = 4 to *k* = 6 and the narrow grey region III allows diversification from *k* = 5 to *k* = 7. For other parameter combinations, only a single diversification event is possible in the infinite NSR model. Limited diversification is one of the major drawbacks of the deterministic SR and NSR models. However, demographic stochasticity may be the key to, at least partially, resolving this puzzle since stochasticity very often leads to different dynamics compared to deterministic models.

**Figure 3:**
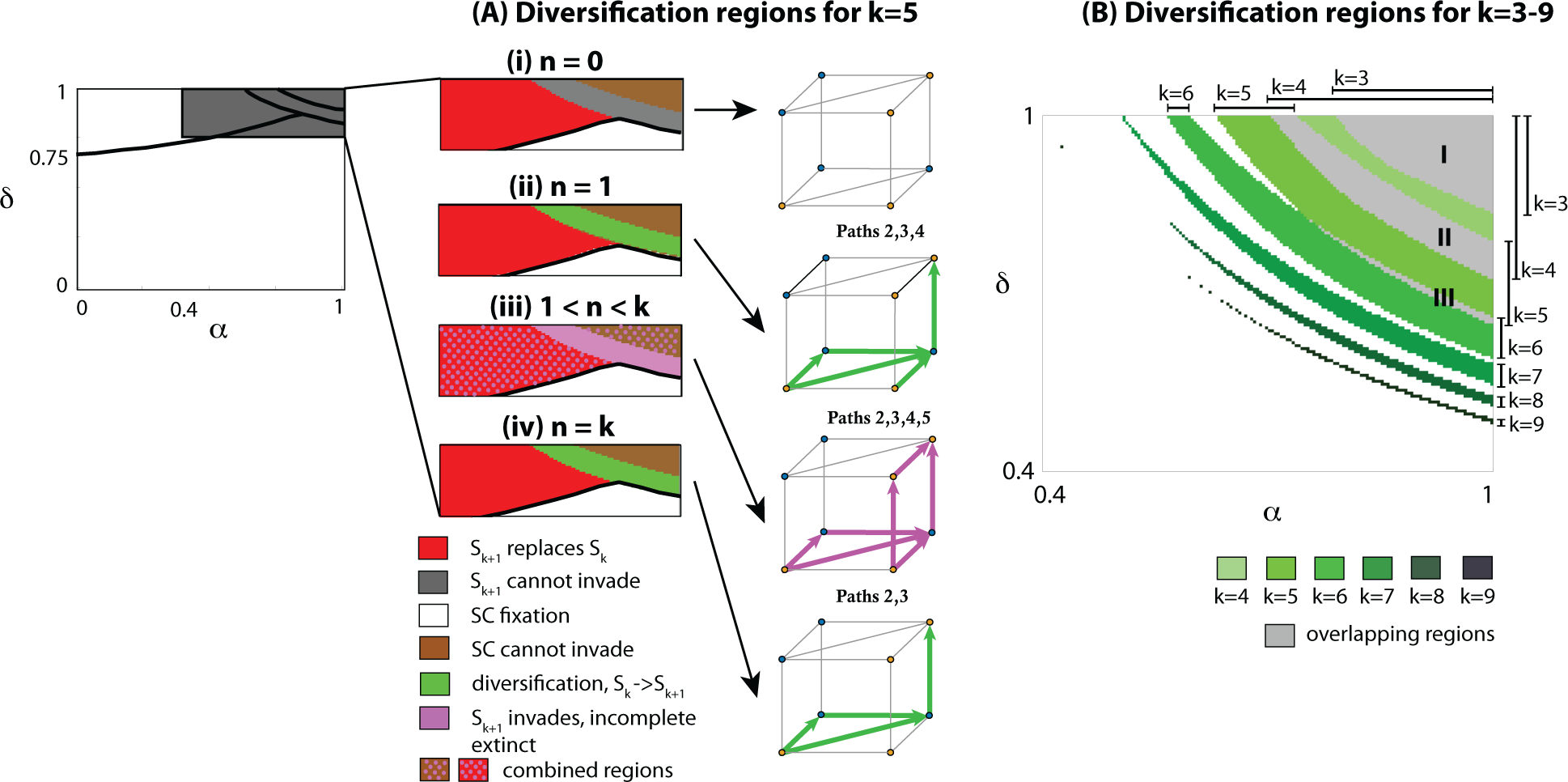
Diversification in the infinite population model. (A) Summary for all pathways 1 - 5 as a function of inbreeding depression (δ) and proportion of self-pollen (α) when the initial number of SI haplotypes (*k*) is five and the level of completeness ranges from zero to k (0 ≤ *n* ≤ *k*;*n* = number of complete haplotypes). The bifurcation plots are superimposed plots of Figures S3-S5 (Supplemental Information). Color coding: diversification with long-term increase in the number of SI haplotypes and a loss of the intermediate mutant (green), short-term diversification (light magenta), no diversification due to exclusion of the novel SI haplotype *S*_*k*+1_ (grey), no diversification because the novel SI haplotype *S*_*k*+1_ replaces its ancestral SI haplotype (red). Below the thick black line is a parameter region where mutations in the SLFs may lead to a complete SC haplotype class (pathways 2 - 4), which results in the fixation of this SC class and a loss of self-incompatibility from the population. Therefore, diversification is only possible in the region above this line. Long-term stable coexistence after a diversification event is possible only for n =1 and *n* = *k* (green regions). For 1 < *n* < *k*, after the invasion of *S*_*k*+1_, all incomplete haplotype classes slowly go extinct. An incomplete SI class can be rescued if it gains the missing haplotype before extinction, and diversification happens if all incomplete classes gain the missing SLF. We highlight all feasible pathways for the particular n in the corresponding cube, using the color of the region where diversification via this pathway is possible either a long-term coexistence of *k* + 1 SI haplotypes (green), or short-term coexistence (light magenta). For *k* ≥ 7 diversification via pathways 2,3,4 for *n* =1 is not possible (Appendix A). For 1 < *n* < *k*, the red and brown regions overlap with magenta for Pathway 5 (light magenta dots). In this region we expect path 5 to dominate the dynamics as in the brown and red regions the number of SI classes remain intact, but pathway 5 modifies the number of SI classes. (B) Diversification regions as a function of the number of SI classes k. These results are identical to Gervais *et al*. (2011) (Figure 2). Regions get smaller as k gets larger and shift to smaller values of α and δ. There is overlap between diversification regions for *k* = 3 and *k* = 4 (grey region I), *k* = 4 and *k* = 5 (grey region II) and *k* = 5 and *k* = 6 (grey region III).

**Figure 4:**
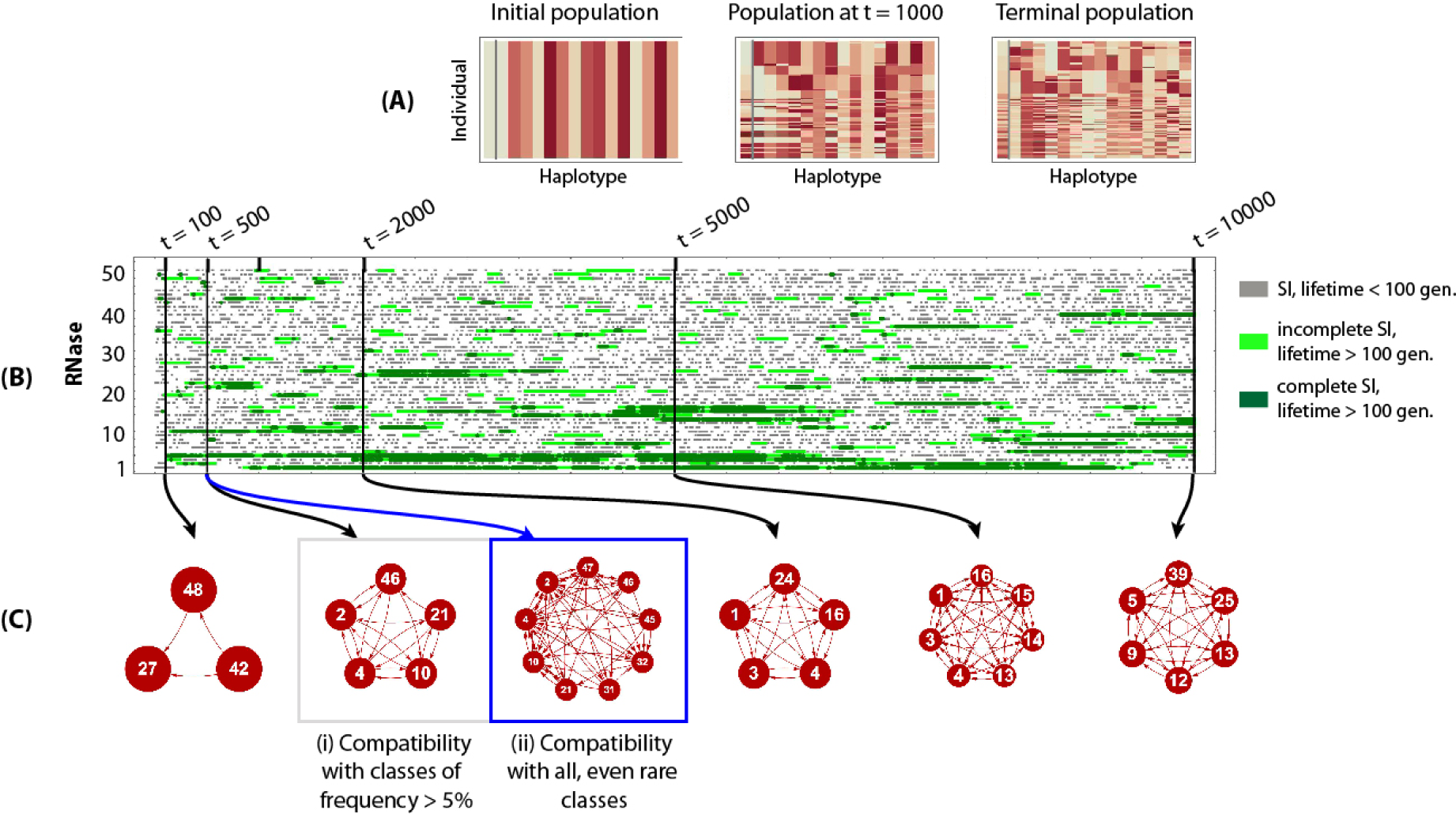
An example of the evolutionary dynamics of the model used for the stochastic simulations with *N* = 1000 individuals, *n*_*R*_ = *n*_*F*_ = 50, *μ_R_ = μ_F_* = 0.001, *α* = 0.8 and *δ* = 0.95. Panel A visualizes the haplotype sequences at three time periods where the first column represents the SRNase and each subsequent column an SLF (there are 2000 rows in each table). In the initial population all haplotypes are self-compatible, with the same SRNase class and the same random set of SLF genes. At t = 1000 and in the terminal population (t = 10000), there are a number of different SRNases and haplotypes can have different sets of SLFs. Panel B shows the appearance/disappearance of the 50 potential self-incompatible (SI) haplotypes classes in the population over 10000 generations. The length of the line represents the lifespan of the haplotype. Grey lines are the short life-span haplotypes (complete or incomplete) that exist for less than 100 generations; light green lines are the incomplete SI haplotypes, while dark green lines are the complete SI haplotypes, which exist for more than 100 generations. Complete haplotypes have all the SLF genes required for all SRNases in the population, while incomplete haplotypes are missing some SLF genes. Panel C shows the compatibility among SI haplotypes in the population at five time periods. Red lines joining haplotypes indicate compatibility, while the absence of a line between haplotypes indicates incompatibility. In all cases except (ii) these are for haplotypes with a frequency > 5%; (ii) shows compatibility among haplotypes including the rare classes. Comparing (i) and (ii) at *t* = 500 shows the differences in mate availability when comparing just the higher frequency (*>* 5%) classes, to when all haplotype classes are included.

### 3.2 Stochastic simulations: an introductory example

The deterministic model for infinite population size appears inadequate to explain the generation and maintenance of the large number of haplotypes observed in natural populations. Firstly, for both SR and NSR systems, the deterministic model predicts that the number of SI haplotypes increases by at most two when selfing rate α and inbreeding depression δ are constant (see also Gervais *et al*. (2011)). Secondly, in our determinstic NSR model, diversification often leads to the eventual loss of all incomplete haplotypes. Consequently, we next examine stochastic simulations, which include features such as unconstrained mutational order and drift. We find that stochastic simulations solve some of the problems of the deterministic model in explaining haplotype diversification.

We begin by presenting an introductory example that clarifies the essential concepts and parameters used in the following sections. All simulations begin (*t* = 0) with a population of self-compatible individuals, with the same SRNase class and a random set of SLF genes including the SLF that corresponds to its own SRNase (Figure 4A). In this example, there are 50 possible SRNase haplotype classes. Only SI haplotypes are presented in (Figure 4B) and these can be classified into two classes: complete haplotypes with all SLF genes corresponding to other SRNase haplotypes in the population, and incomplete SI haplotypes that are missing some SLFs. Simulations are run for 10000 generations and the dashed line for each SRNase class represents the emergence of that class; grey, short lines are SI haplotypes that have a lifetime of less than 100 generations, light green lines are incomplete haplotypes with a lifespan of greater than 100 generations and dark green complete haplotypes with long lifespan (¿100 generations, Figure 4B). The length of the line shows lifespan and so we see that, in general, complete haplotype classes tend to have a longer lifespan than incomplete haplotypes (Figure 4B). Completeness (having a full set of SLF genes) determines mate availability. In this example, at generation 500, haplotype class 4 is complete and can therefore mate with all other haplotypes (Figure 4Ci,ii). In comparison, haplotype class 21 is incomplete and has reduced mate availability, both when considering higher frequency (> 5%, Figure 4Ci) and rare classes (Figure 4Cii). At all time periods, the higher frequency classes (> 5%) tend to be complete and able to mate with all other higher frequency classes in the population (Figure 4C).

### 3.3 Stochastic simulations: Establishment and the number of SI classes

Next, we examine the conditions associated with S haplotype diversification. We found that the frequency of SI haplotypes was greatest (> 75%) at intermediate to high values of self-pollen deposition (α = 0.6 - 1) and high inbreeding depression (δ > 0.85) (Figure 5a). Here, the average frequency of SI types in the population was highest (closest to 1) with high values of both self-pollen deposition (α > 0.8) and inbreeding depression (δ > 0.86) (Figure 5b). In the parameter space (α > 0.4, δ > 0.82), the average number of SI haplotypes that evolved was 7-14 for population size 1000 (Figure 5b), although some of these are rare. Population size influenced the relative frequencies of SI to SC haplotypes, so that the overall relative frequency of SI to SC increased with population size (see the results for *N* = 200 in the Supplemental Information S1).

**Figure 5:**
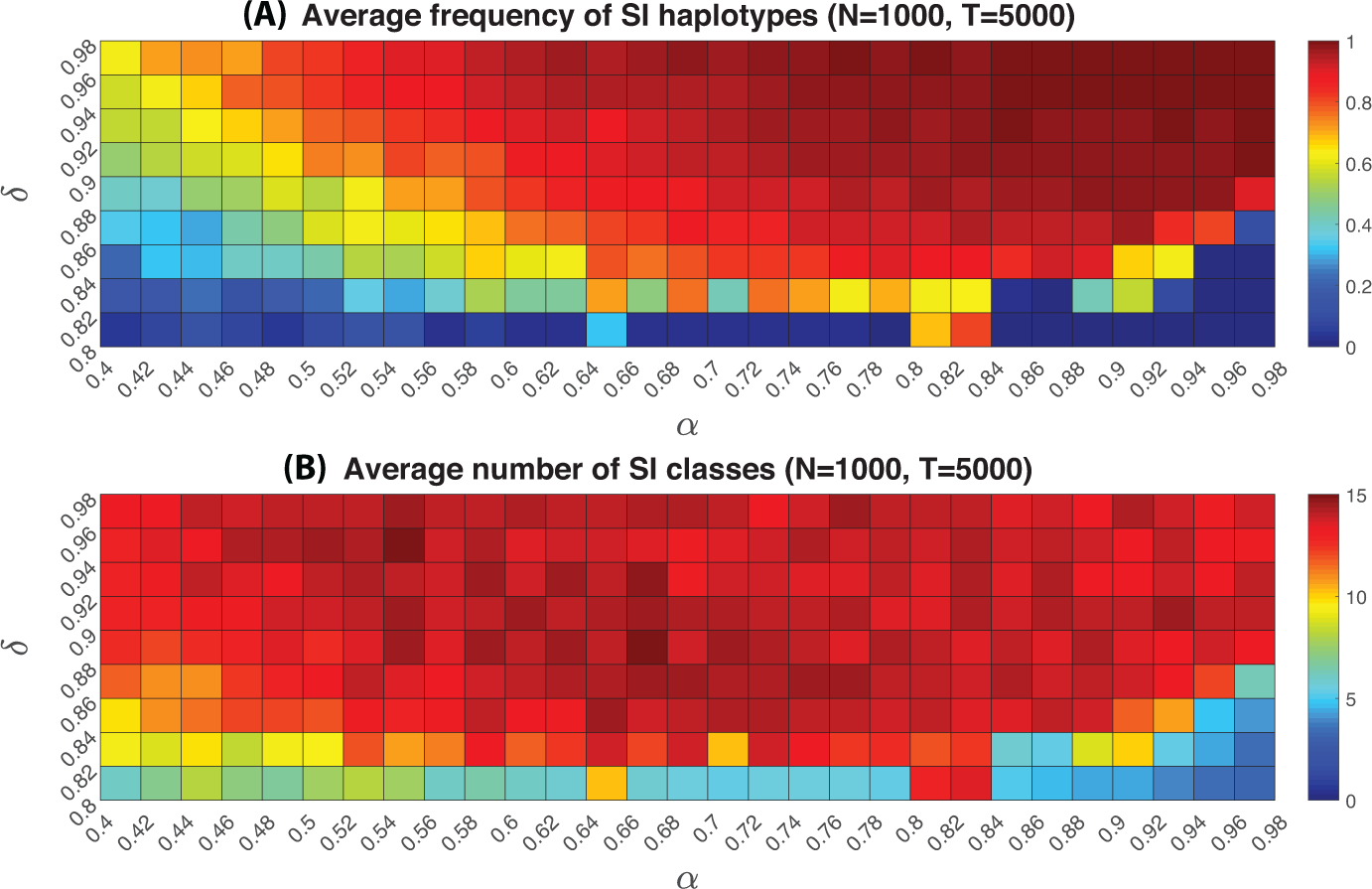
The average number (a) and frequency (b) of self-incompatibility types in a non-self recognition system in a finite population (*N* = 1000) as a function of self-pollination rate (α) and inbreeding depression (δ). Colours in the grid squares represent a gradient from low (green) to high (red) average numbers of SI types (a) and total frequency of SI types (b) in the population. Mutation rate *μ*_*R*_ = *μ*_*F*_ = 0.001, Length of the F-box (SLF) sequence *L* = 15, and the total number of potential SLF and SRNase types *n*_*F*_ = *n*_*R*_ = 50. Analogous plots for small population size *N* = 200 are presented in the Supplemental Information, Figure S1.

### 3.4 Stochastic simulations: evolutionary dynamics and the interplay between SI and SC classes

Our model shows the three stages in the evolution of a NSR SI system: the establishment, diversification and stationary phases (Figure 6). During the establishment phase the relative frequency of SC haplotypes decreases as these are replaced by SI haplotypes. Once the SI system is established (defined as *k* = 3), the population enters the diversification and stationary phases. During the stationary phase, there were many cases of equal frequencies of SI classes, but also some low frequency SI classes and the recurrence of SC haplotypes. The occurrence of SC haplotypes was more likely when the mutation rate was higher (*μ* = 0.001) and when there was a larger number of potential haplotypes (*n*_*R*_ = *n*_*F*_ = 50) (Figure 6C). A rapid change in SC and SI haplotype frequencies was most prominent at lower mutation rates (Figure 6B and D), independent of the number of potential haplotypes (*n*_*R*_ and *n*_*F*_). For higher mutation rates (Figure 6a and c) the frequency of SC haplotypes was lower and these were more transient, but their occurrence increased with larger potential haplotype number (*n*_*R*_ = *n*_*F*_ = 50; Figure 6C). These patterns were qualitatively consistent for a and δ combinations within the parameter space where SI invades (see Figure 5). However, we found that the frequency of SI types was greater for the parameter space where new SI haplotypes are more likely to invade. Equal frequencies of the most common SI classes were apparent in the stationary phase, and this was most consistent for lower mutation rates (Figure 6B and D). Low frequency SI haplotype classes were, however, also present at sufficient numbers leading to the occasional replacement of the most common SI classes by the low frequency SI classes.

**Figure 6:**
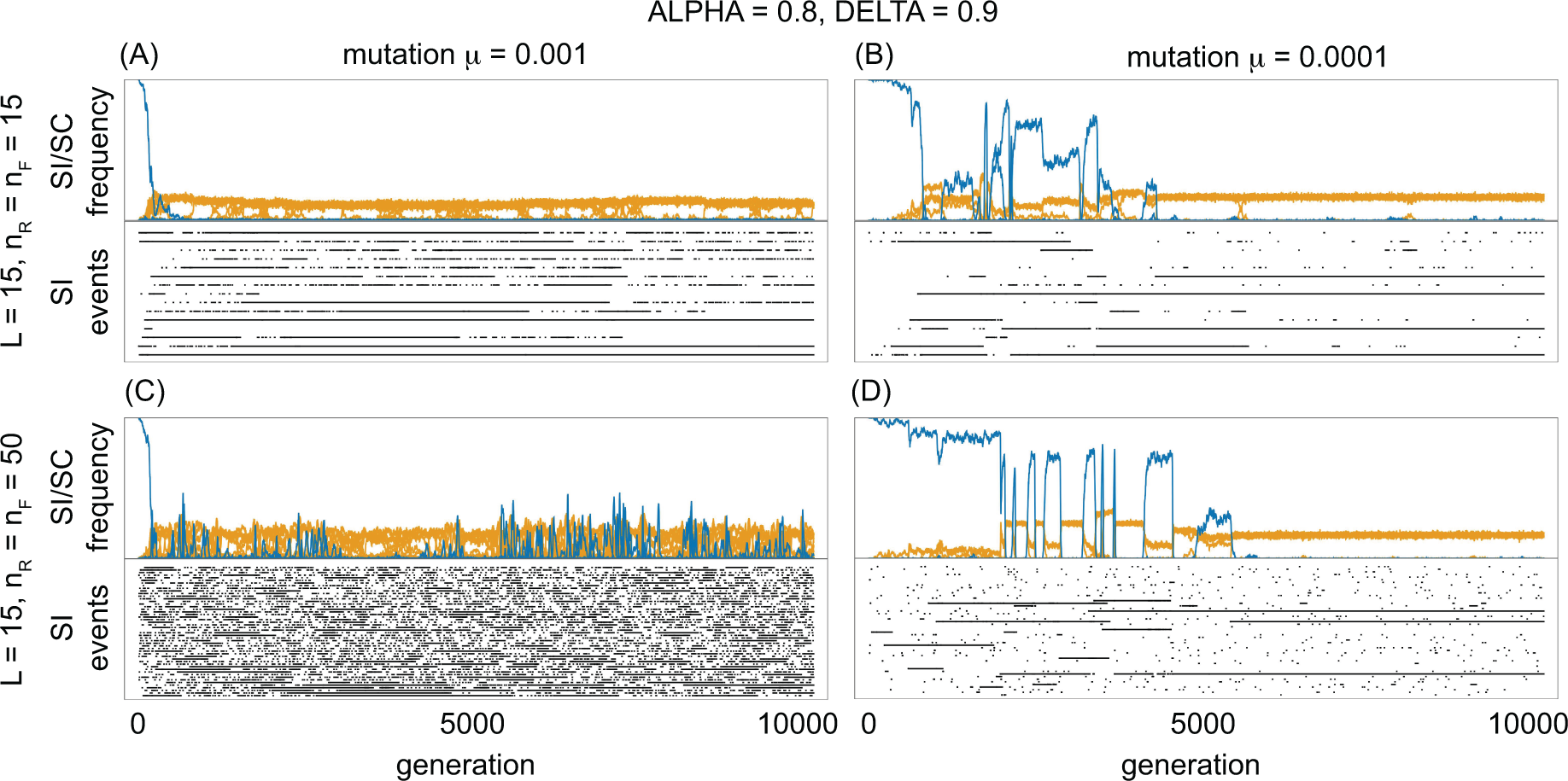
The evolutionary dynamics of the non-self recognition self-incompatibility model in relation to mutation rate and number of possible SNRase (*n*_*R*_) and SLFs (*n*_*F*_). For each panel (A to D) the upper graph represents the frequency of self-incompatible (SI, yellow) and self-compatible (SC, blue) haplotypes (frequencies sum up to one) over 10000 generations and the lower graph is the occurrence of events for SI haplotypes. Each line in the lower graph shows an event that results in a new SI haplotype class arising from a population without the given SRNase. This may arise from either a SC or SI haplotype (see Figure 2). For A and B the number of potential SRNase and SLFs *n*_*R*_ = *n*_*F*_ = 15, while for C and D the number of potential SRNase and SLFs *n*_*R*_ = *n*_*F*_ = 50. For A and C mutation rate *μ*_*R*_ = *μ*_*F*_ = 0.001, while B and D mutation rate *μ*_*R*_ = *μ*_*F*_ = 0.0001. For all models, *L* + 1 is the length of the haplotype (one SRNase plus the L = 15 spaces for SLFs). Simulations were run at values of self-pollination rate α = 0.8 and inbreeding depression δ = 0.9.

To track SI haplotypes (ignoring SC haplotypes), we recorded the first occurrence of a given class (Figure 6) that may have arisen from either a SC or SI haplotype (see Figure 2). Each line (row) therefore shows an event that begins with the occurrence of a novel SI haplotype and ends with the extinction of the last SI haplotype from that class; the length of the line is therefore the lifespan of the haplotype class. Short events occur when the haplotype class is lost due to demographic stochasticity, while the long events represent haplotype classes that reach sufficient frequency and are maintained in the population. There were many more short than long events and the lifespan of SI haplotype classes varied in relation to mutation rate and potential haplotype number. Higher mutation rates led to a greater proportion of short events (Figure 6A and C). Moreover, for a given mutation rate, there was higher turnover and less stability of the SI classes when there were more potential haplotypes (*n*_*R*_ = *n*_*F*_ = 50; Figure 6A and C). Consequently, the highest turnover of SI classes occurred at higher mutation rates and numbers of potential haplotypes (Figure 6C), compared to the longer lifespans and less turnover observed for lower mutation rates and number of potential haplotypes (Figure 6B).

**Figure 7:**
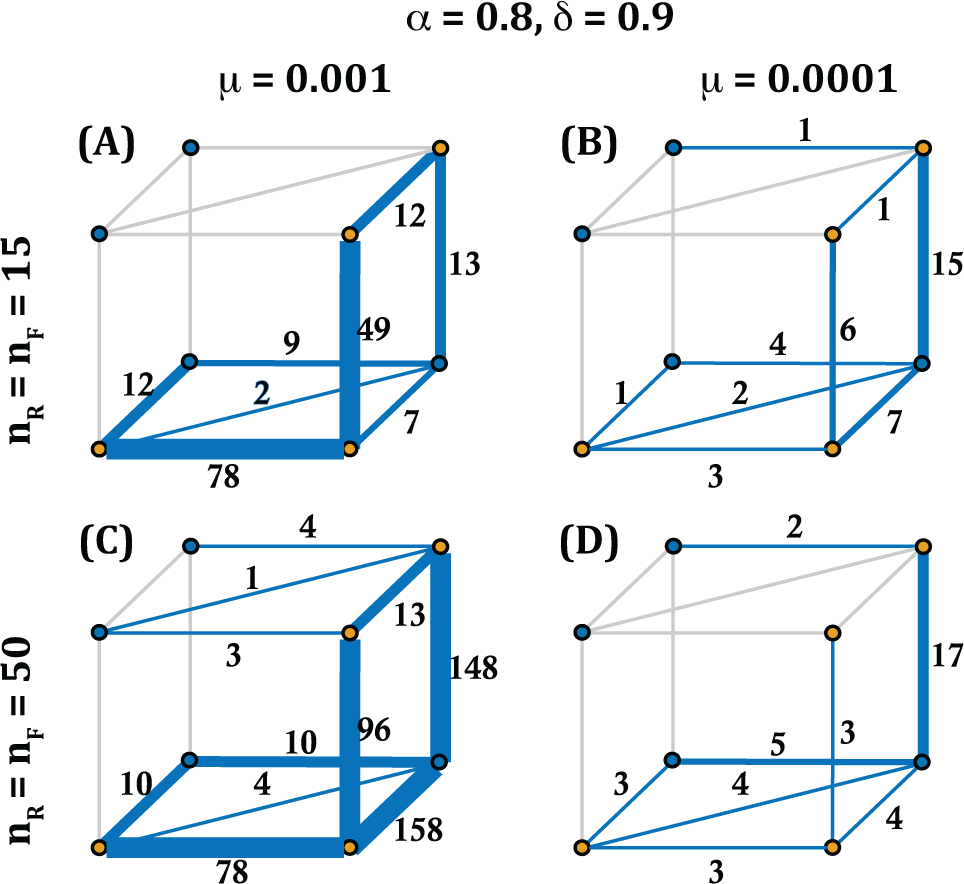
The likelihood of transitions along different evolutionary pathways for S haplotypes with long life spans as a function of potential haplotype number (*n*_*R*_ = *n*_*F*_) and mutation rate. Long lifespan haplotypes were those maintained in the population for more than 100 generations. For (a) and (b) the number of potential haplotypes is 15 (*n*_*R*_ = *n*_*F*_ = 15), and (c) and (d) *n*_*R*_ = *n*_*F*_ = 50. For (a) and (c) mutation rate = 0.001, while for (b) and (d) the mutation rate = 0.0001. The numbers on each side of the cube represent the number of transitions between two vertices. Each vertex represents the states outlined in Figure 2A, with blue vertices for self-compatible and yellow for self-incompatible haplotypes. Simulations were run with the following parameter values: *N* = 1000, self-pollination rate α = 0.8 and inbreeding depression δ = 0.9.

### 3.5 Stochastic simulations: Most likely evolutionary pathways for SI haplotypes with long lifespans

We now examine whether there is an association between the evolutionary pathway of long lifespan SI haplotypes for different mutation rates (*μ*) and potential haplotype numbers (*n*_*R*_ = *n*_*F*_). Both mutation rate and potential haplotype number influenced the likelihood of each pathway for SI haplotypes with a lifespan of more than 100 generations. High mutation rate (*μ* = 0.001) and a lower number of possible haplotypes (*n*_*R*_ = *n*_*F*_ = 15) favored the pathway that maintains self-incompatibility (Figure 7A). Yet, with a greater number of possible haplotypes (*n*_*R*_ = *n*_*F*_ = 50), a higher mutation rate resulted in high transition likelihoods for both pathway 4 with a SC intermediate and for pathway 5 that maintains SI (Figure 7C). In comparison, for lower mutation rates (*μ* = 0.0001), we observed a larger number of transitions through the pathway with an SC intermediate (pathway 4), and this was more consistent for different potential haplotype numbers (Figure 7B, D). Furthermore, and aligned with our predictions on the unlikeliness of pathways 1 and 2 (they require a simultaneous loss and gain of two specific SLF), we don’t observe these pathways in our simulations (see above for more discussion on pathway 1).

### The effect of completeness

Complete haplotypes with no missing SLF genes had the longest lifespan and maintained the highest frequencies compared to haplotypes missing between two and four SLF genes (minimal completeness deficit between two and four; Figure 8). In comparison, missing only one SLF gene had a smaller effect on lifespan and frequency. All completeness deficits were represented in the short lifespan and low frequency classes. However, the complete haplotypes, or those missing only one SLF gene, had the longest lifespans (> 100 generations) and highest (> 0.1) frequencies (Figure 8). We provide further illustration of the importance of completeness for haplotype lifespan in the Supplemental Information, Figure S2. In this example, we present a single representative SI class and its completeness deficit throughout its lifetime showing that extinction of an SI class is associated with the loss of completeness.

## 4 Discussion

We use analytical theory and stochastic simulations to examine the conditions under which novel S haplotypes evolve for non-self recognition (NSR) self-incompatibility. In addition to a pathway that maintains self-incompatibility, we found that diversification may occur through self-compatible intermediates. The parameter space for diversification, and whether it resulted in long- or short-term diversification, also varied with evolutionary pathway and with both the level of inbreeding and self-pollination. Our results also highlight the importance of completeness and haplotype number for diversification in NSR systems, how this varies with evolutionary pathway and how this interaction determines the long-or short-term coexistence of novel haplotypes. For finite populations (N = 200-1000), we found that 7-14 S-haplotypes evolved under conditions of high inbreeding and moderate to high self-pollination, while the frequency of SI types was close to one when both self-pollination and inbreeding depression were high. For infinite populations, the number of S-haplotypes increased by at most two S-haplotypes even though, in principle, the number of S-haplotypes can increase to infinity. Moreover, the parameter space for diversification was considerably smaller for infinite than finite populations. Mutation rate and number of possible haplotypes influenced the evolutionary dynamics of the NSR model: overall, higher mutation rate and greater number of potential haplotypes (*n*_*R*_, *n*_*F*_, *L*) resulted in shorter lifespans and a greater turnover of SI classes. Yet, when considering long lifespan haplotypes, these conditions influenced the probability of evolving through a pathway that maintained SI or one with a SC intermediate.

**Figure 8:**
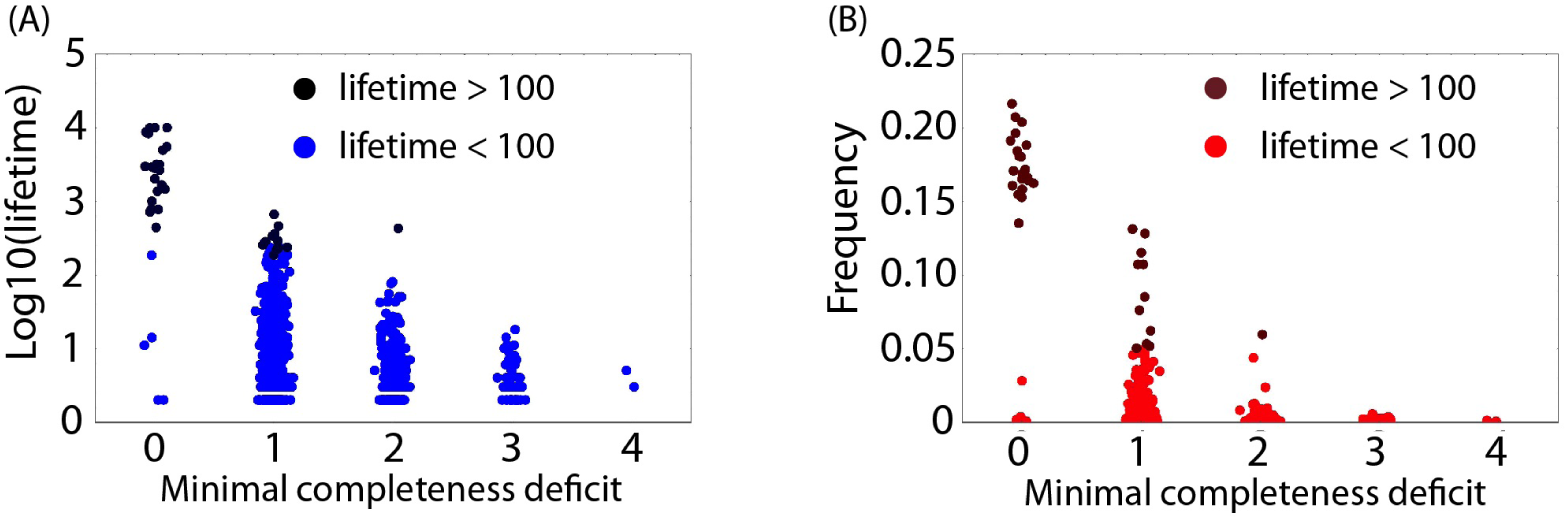
Completeness deficit of the S-alleles. (A-B) Self-incompatible (SI) haplotype lifespan and frequency in relation to its minimal completeness deficit. The light/dark colors represent short/long events (lifetime greater/shorter than 100 generations). Completeness deficit is a measure of how many SLF genes a haplotype is missing that relate to potential mating partners in the population. Minimal completeness refers to the minimum through all haplotypes in that SI class over the entire lifespan. This measure is therefore related to mate availability so that a minimal completeness deficit of 0 is when the haplotype has the full set of SLF genes that are able to detoxify all potential SRNases in the population (i.e., full mate availability); A minimal completeness deficit of 1 implies that the haplotype is missing one SLF associated with an SRNase in the population and is therefore unable to mate with individuals with this SRNase. Consequently, the higher the minimal completeness deficit, the lower the mate availability for that haplotype. Simulations were run with the following parameter values: *N =* 1000, *n*_*R*_ = *n*_*F*_ = 15, *L =* 15, *μ*_*R*_ = *μ*_*F*_ = 0.001, *α =* 0.6 and *δ =* 0.9. For the purpose of clarity an additional scatter was introduced to the completeness deficit data.

We first discuss the conditions that may promote diversification and how these may vary through the evolutionary process. Then, we relate our results to previous models and examine how they extend our current understanding of novel S haplotype evolution in NSR systems. We conclude by discussing future directions for theoretical models and how these, when combined with genomic data, can provide insight into the evolution of diversity in NSR systems.

### 4.1 Evolutionary pathways for diversification: self-pollination, inbreeding depression and haplotype number

Our results show that self-pollination and inbreeding depression influenced diversification outcomes; yet this varied for the different evolutionary pathways and with different levels of completeness and haplotype number. We first discuss how these parameters influence diversification for each pathway and then discuss the interaction between selfing rate and inbreeding depression

#### First mutation in the female-specificity with a SC intermediate

We found that diversification was unlikely through a pathway where the first mutation happens in the SRNase resulting in an intermediate SC haplotype (pathway 1); this pathway is identical to the path presented in Uyenoyama *et al*. (2001) and Gervais *et al*. (2011). Although this pathway is in principle possible (see Figure 3), it is unlikely to contribute to the observed diversity because, when comparing all pathways simultaneously, this parameter region overlaps with the region where SI is lost from the population via alternative pathways (region below the thick line in Figure. 3). Furthermore, this pathway is not observed in the simulations. This discrepancy likely originates from differences in the order of mutations between the two theoretical approaches. In contrast to the analytical model, where diversification through a pathway occurs in a set sequence, the order of mutations is random in simulations. This means that, like Gervais *et al*. (2011), pathway one was not observed in our simulation results because either an SLF or SRNase mutation can occur first resulting in the loss of the SI system. Consequently, although theoretically possible, pathway 1 can only occur under the restricted conditions observed in the deterministic model and is unlikely to contribute to long-term diversification.

#### First mutation in the male-specificity with a SC intermediate

Haplotype number influences the strength of balancing selection during S allele diversification. In our analytical model, long-term diversification was more likely with lower haplotype numbers for pathways where the first mutation happened in the male-specificity (e.g. Figure 3). This is because for lower haplotype numbers the intermediate SC haplotypes have an advantage over SI haplotypes due to self-pollination and thus can invade the population, eventually resulting in diversification after the next mutation. This effect gets weaker with increasing numbers of haplotypes as SI haplotypes become less mate-limited, resulting in a smaller parameter region for diversification and this occurs at lower inbreeding depression. Consequently, there is generally no overlap in the parameter space for diversification for different numbers of haplotypes initially present in the population. Thus, surprisingly, even though in principle the number of S-haplotypes can increase to infinity in our analytical infinite population model, the number of haplotypes increases by at most two for any fixed level of inbreeding depression and self-pollination. However, in nature, variation in selfing rate and inbreeding depression may facilitate traversing non-overlapping parameter spaces. An equivalent result was found for self-recognition systems in Gervais *et al*. (2011), indicating the importance of frequency-dependent selection for evolutionary outcomes. This has interesting implications for considering the application of these models to empirical data on the distribution and number of S haplotypes (see Discussion below). For example, small populations with fewer S alleles may provide the conditions that promote diversification in a metapopulation.

#### First mutation in the female-specificity with a SI intermediate

In contrast to the above pathways where diversification was unlikely for higher haplotype numbers, diversification for a pathway where SI is maintained (pathway 5; Figure. 3) is possible for any initial haplotype number. Moreover, since all haplotypes outcross, this pathway has no fitness costs associated with self-pollination and inbreeding depression. This implies that, in principle, multiple consecutive diversification events are possible for this pathway for any level of inbreeding depression and self pollination. However, in our deterministic model the diversification was only short term as all incomplete SI haplotypes slowly go extinct.

#### Purging of deleterious alleles

Theory predicts a negative relationship between inbreeding depression and selfing rate, such that the purging of deleterious recessive alleles through selfing reduces inbreeding depression (Lande and Schemske 1985). However, the combination of high inbreeding depression and selfing rate is not unrealistic, given that studies have found inbreeding depression in populations and species with high self-fertilization rates (Byers and Waller 1999; Winn *et al*. 2011). Moreover, Gervais *et al*. (2014) found that purging had little effect on the spread of self-compatible individuals if most deleterious alleles had weak fitness effects. We found that the combination of self-pollination and inbreeding depression where diversification was observed varied with evolutionary pathway. Indicating the potential for different conditions to favour diversification through alternative pathways. In our model, inbreeding depression was fixed through time (i.e. purging was not considered), even though its strength may vary with population size (Bataillon and Kirkpatrick 2000) and the degree of bi-parental inbreeding (Porcher and Lande 2016). Sheltered genetic load may also influence the dynamics of deleterious alleles and inbreeding depression (Porcher and Lande 2005; Llaurens *et al*. 2009), although this is more likely to be important for sporophytic SI systems with dominance hierarchies among alleles (Llaurens *et al*. 2009). By considering the importance of dynamic inbreeding and genetic linkage, future models could further examine how these apply to the evolution of novel S haplotypes in an NSR SI system.

### 4.2 Evolutionary pathways for diversification: the effect of completeness

The relative importance of completeness for diversification may vary for self-vs. non-self recognition systems. Studying an SR model, Sakai (2016) found that an incomplete SI system was essential for diversification, and that this occurred during the initial evolution of the SI system. Here the pollen genes (male component) for a given specificity were not fully rejected by the female-determining genes for that specificity: leading to incomplete rejection following the matching of SI haplotypes. Sakai (2016) interprets this model of diversification during SI establishment as reflecting the transpecific sharing of S alleles, where S alleles evolved before the species split (Sakai 2016). In our model, incompleteness (missing SLF genes related to SRNases in the population) reduces mate availability and fitness. One of the key results of this study on NSR SI is the importance of completeness for long-term diversification. Long-term diversification was only observed when one haplotype is complete (*n* = 1) or all haplotypes are complete (*n* = k). For all other completeness levels only short-term diversification is possible because incomplete haplotypes slowly decrease in frequency due to their reduced fitness (see the exact analytical conditions in the Appendix A). Here, incomplete haplotypes slowly go extinct, unless rescued by mutations that restore completeness (a full set of SLFs). This is analogous to evolutionary rescue (Gomulkiewicz and Holt 1995; Gonzalez *et al*. 2013), where mutation can prevent extinction, enabling the haplotype to persist in the population. Our simulation data also show the potential importance of completeness for haplotype lifespan and turnover. We found higher turnover of SI haplotypes when the number of potential haplotypes was larger. Here, when there were more potential haplotypes (*n*_*R*_ = *n*_*F*_ = 50), a larger proportion of the SI haplotypes had a higher completeness deficit (i.e. more missing SLFs). Given that mate availability scales with completeness deficit, incomplete haplotypes with more missing SLFs are likely to be selected against, reducing their lifespan and contribution to long-term diversification outcomes.

#### Completeness and the pathway that maintains SI

Our results also raise a number of questions about diversification via the pathway that maintains SI (pathway 5). Diversification is not possible for this pathway when the number of incomplete haplotypes is zero or one (i.e. *n* = 0 or 1). When the number of complete haplotypes is greater than one diversification is possible, but only if all incomplete haplotypes rapidly become complete, otherwise they go extinct. This challenges the feasibility of the evolutionary pathway for diversification proposed by Fujii *et al*. (2016). Yet, when we consider finite populations we do see diversification through this pathway. This implies that conditions in the simulations such as a random order of mutation events, higher mutation rates, finite population size and having a more flexible SLF template (i.e. more SLFs to begin with) may facilitate diversification through this pathway. In conclusion, comparing the results of our deterministic and stochastic models highlights the potential importance of completeness for long-term diversification in the NSR SI system. These results, however, are based on the assumption of global dispersal of pollen. It is possible that the importance of incompleteness may decrease if pollen has a limited range, as this may reduce the effects of missing SLFs on mate availability. Future models that extend our results to non-global dispersal may therefore assist our understanding of the role of incompleteness in S haplotype diversification.

### 4.3 Congruence of theoretical models and empirical data: estimates of haplotype number

The extremely high level of S haplotype diversity observed in natural populations is well established (Lawrence 2000; Castric and Vekemans 2004), raising interesting questions about the congruence of theoretical models with empirical data. We found that the number of S haplotypes derived from our model (*k* = 7 - 14, for population size 1000) were far fewer than the diversity commonly observed in natural populations of species with SR and NSR SI (20-40 SI haplotypes; Lawrence (2000)). These results are similar to the simulation results of Gervais *et al*. (2011) who found that diversification peaked at between 7-18 alleles. It has been suggested that population structure may provide the conditions for diversification (Uyenoyama *et al*. 2001; Gervais *et al*. 2011). Incomplete reproductive barriers and hybridization among species may also create the population structure required to enhance diversification (Castric *et al*. 2008). In this case, novel S-haplotypes may evolve in separate species, which are then exchanged among species via introgression. These novel S alleles may spread through the population once they are decoupled from the hybrid genetic background. These ideas are intriguing given the transpecific nature of S alleles and the maintenance of SI during speciation (Igic *et al*. 2004). Further models that include population structure, as well as introgression, are therefore required to assess the potential importance of this for diversification at the S locus. The total number of S haplotypes predicted by Sakai (2016) was higher (40-50 alleles), and more in line with population estimates. The mechanism of diversification in this model, however, suggests that novel S alleles evolve during the initial evolution of the SI system from self-compatibility. It is also based on a self-recognition SI, and involves variation in the strength of the incompatibility reaction, suggesting that the mechanism may be less applicable to S haplotype diversification in NSR systems.

There are a number of assumptions required to simplify the inherent complexities of modeling the evolution of new S-haplotypes in NSR systems. These assumptions may have contributed to the lower haplotype estimates compared to that observed in natural populations. We firstly assume that there is no recombination within the S locus. However, Kubo *et al*. (2015) provide evidence of genetic exchange at the S-locus for Petunia and suggest that SLFs may be shared among S-haplotypes via this process. Inclusion of genetic exchange may therefore facilitate novel S-haplotype evolution, as suggested by Fujii *et al*. (2016) who found that this had a important role in evolution of novel S-haplotypes in their NSR model. Secondly, we assume equal mutation rates for male- and female-determining components of the S locus. Given the larger size of the SLF compared to SRNase genomic region, and some evidence of greater variation and turnover of SLFs (Kubo *et al*. 2015), the potential influence of higher mutation rates for SLF genes could be tested. Consequently, extending our model to include genetic exchange, unequal crossing over and variation in mutation rates for male- and female-determining components may result in different evolutionary outcomes and equilibrium number of S-haplotypes.

### 4.4 Haplotype lifespan and frequency: mate availability and negative frequency-dependence

The interaction between haplotype completeness and mate availability can influence the lifespan and frequency of novel SI haplotypes. In the NSR SI system modeled here, mate availability is inversely related to the deficit in SLF genes (minimal completeness deficit), so that complete haplotypes, with no missing SLFs, have the highest mate availability and are able to mate with all other haplotypes in the population. Our results of longer lifespan and high frequency for complete haplotypes (minimal completeness deficit of zero), reflects its importance to mate availability and fitness. Interestingly, and in contrast to our infinite population model, the moderate to high longevity and frequency of haplotypes missing one SLF (minimal completeness deficit of one) suggests that these haplotypes can still maintain high fitness. The deficit in SLF genes may also affect the strength of negative frequency dependent selection (NFDS), so that NFDS weakens when there are fewer mating partners. This may contribute to the stochastic loss of haplotypes with a greater deficit in SLF genes, which is reflected in their shorter lifespans and lower frequencies. Taken together, these results highlight the importance of NFDS and suggest that both evolutionary pathway and mate availability contribute to the outcomes and success of novel SI haplotypes. It would be interesting to see if these results are still apparent with extensions to the model to include local pollen dispersal, since this may lessen restrictions in mate availability.

## 5 Conclusions

Our model demonstrates that novel S haplotypes can evolve across a range of parameter values (inbreeding depression and self-pollination), but that this varies with evolutionary pathway. This result generates intriguing questions about the role of self-compatible intermediates in S allele diversification and how different conditions may favour alternative pathways. For example, when considering empirical data, does the presence of low frequency self-compatible individuals in populations (Raduski *et al*. 2012) represent points in the diversification process? This also raises questions regarding the viability of self-compatible individuals as intermediates for the evolution of new specificities under different models of inbreeding depression. Extensions of this model to include population structure may also help to reconcile differences between theoretical models and the number of alleles commonly observed in plant populations. Combining genomic data with model predictions could provide insight into the evolutionary dynamics of NSR self-incompatibility. For example, variation among individuals in SLF gene position and copy number may provide information on recombination frequency and gene duplication events (see Kubo *et al*. (2015)); while the distribution of mutations within SLF genes may indicate the steps required to produce a novel SLF during allelic diversification. Indeed, previous studies have provided some estimates of the number of mutations required to alter S allele specificities (Matton *et al*. 1999; Chookajorn *et al*. 2004), but given differences in the molecular mechanism and variation in size of the female- and male-determining components, this may vary with SI system. Combining theoretical models with data on the genomic structure of the SLF region will therefore improve our understanding of the diversification process in NSR system: providing a fascinating example of the evolutionary dynamics involved in genetically based recognition systems.

## 6 Acknowledgements

We thank Deborah Charlesworth for helpful comments on the manuscript. The research leading to these results has received funding from the European Union’s Seventh Framework Programme (FP7/2007-2013) under grant agreement number 329960, ERC research agreement number 250152 and REA grant agreement number 291734. We also thank Vincenzo Natali for his influential movie The Cube that was a great source of inspiration. Here, the characters move through cube-shaped rooms with various death traps, as do the S-alleles in our work, which seem to be searching for a successful escape route through a sequence of mutational cubes, facing the self-compatibility and incompleteness traps.

## A The coexistence of complete and incomplete SI haplotypes and the necessary condition for diversification

In this section we are interested in whether SI haplotypes, some complete and some incomplete (missing one SLF), can be stably maintained in the population. Then, we discuss how the derived conditions can be used to study the feasibility of the diversification pathways 1 to 5 discussed in the main text.

In this study, because we aim to examine the conditions associated with each diversification pathway, we study the coexistence of the ”final” number of (complete and incomplete) SI classes after a diversification event. That is, we construct a set of equations, using (3), to describe the dynamics of *n* SI haplotypes *S*_*i*_ that have *F*_*k*+1_, m SI haplotypes *S*_*α*_ that are missing *F*_*k*+1_, as well as the ”novel” haplotype *S*_*k*+1_, where *n* + *m* = *k*. Therefore, and in line with the assumptions on diversification pathways in the main text, we study the possible coexistence of *n* +1 complete haplotypes (*n* complete resident types plus a complete type *S*_*k*+1_) and *k* + 1 - n = m incomplete SI haplotypes which lack the SLF for *S*_*k*+1_. Notice that the haplotype *S*_*k*_ (in the main text used as the ancestral haplotype) belongs to one of the n haplotypes if it has *F*_*k*+1_ (in the main text 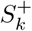), and if it lacks *F*_*k*+1_ it belongs to one of the m haplotypes (in the main text 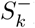)).

For convenience, we will use notation *S*_*i*_ (more generally, we use indices *i, j*,…) to describe all the complete resident haplotypes *S*_1_,…,*S*_*n*_ and call it a group of *n* complete SI haplotype classes. Similarly, *S*_α_ (more generally, we use indices α, β,…) will denote the group of all incomplete SI types. To simplify our notation we write 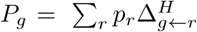 for the sum of all haplotype groups (pollen) that can fertilize a (diploid females) genotype group *g*. This change of notation (and its slight abuse) will help considerably in writing the dynamical model. Similar notation was also used in models of Uyenoyama *et al*. (2001) and Gervais *et al*. (2011).

## A.1 The model

We first write equations for the most general model where *n, m* ≥ 2, implying *k* ≥ 4, after which we discuss the remaining four cases *n* = 0, *m* ≥ 3; *n* =1,*m* ≥ 2; *n* ≥ 3, *m* = 0 and *n* ≥ 2, *m* =1. See the equations and the explanation below for why such a distinction must be made. The expected fitness for all genotype groups (see the definition above), in the most general case (*n, m* ≥ 2), can be written as

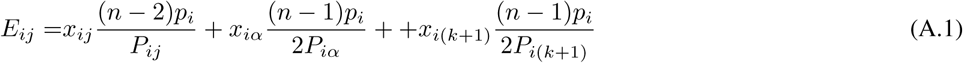

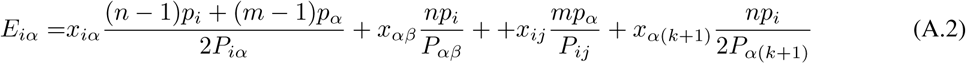

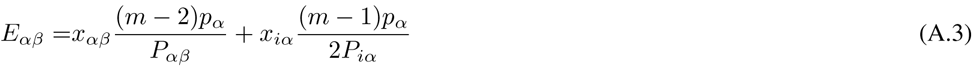

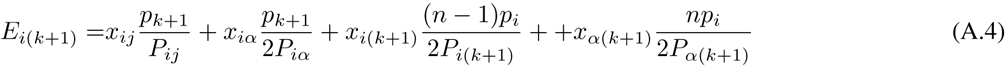

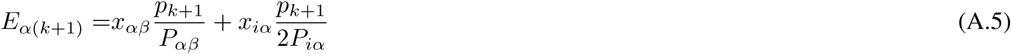

The frequency of all haplotypes that can fertilize can diploid group are

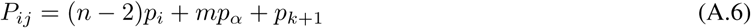

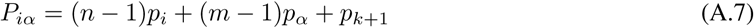

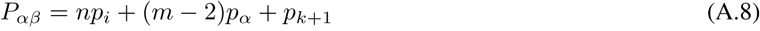

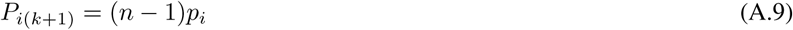

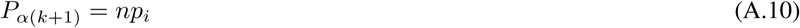

and the haplotype group frequencies in terms of genotype group frequencies are

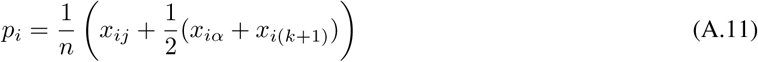

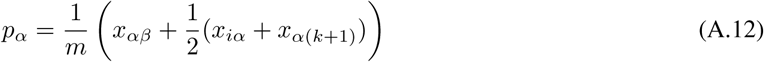

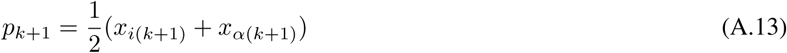

We now examine how these equations were built. For example, *E*_*ij*_ denotes the expected fitness of an arbitrary genotype where both haplotypes belong to the group *S*_*i*_. Therefore, looking at the first term in (A.1), females in group *S_i_S_j_* with frequency *x*_*ij*_ will produce a *S_i_S_j_* offspring with probability 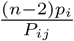. This is because there are *n* - 2 *S*_*i*_ haplotypes that can fertilize an arbitrary diploid female in this group (the frequency is (*n* - 2)*p*_*i*_), and *P*_*ij*_ is the frequency of all the pollen that can fertilize an arbitrary diploid genotype in this genotype group. Similarly, the second term in (A.1) says that females in group *S*_*i*_ *S*_*α*_ with frequency *x*_*iα*_ will be fertilized by haplotypes from group *S*_*i*_ to produce a *S_i_S_j_* offspring with probability 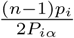. This is because there are *n* – 1 *S*_*i*_ haplotypes that can fertilize an arbitrary diploid female *S_i_S_α_* (the frequency is (*n* - 1)*p*_*i*_), *P*_*iα*_ is the frequency of all the pollen that can fertilize an arbitrary genotype in this diploid genotype group, and then with probability one half the offspring is of type *S*_*i*_*S*_j_.

## A.2 Some general comments

There are a number of reasons why we must write separate models for the special cases *n, m* = 0,1. Firstly, notice that when n, *m* = 0, 1 all expressions *P*_*g*_ will have zero or negative terms because there are not enough haplotype classes in *S*_*i*_ or *S*_*α*_ that can fertilize the corresponding genotype group. This is easily corrected by neglecting all the negative terms (e.g. by multiplying the term by an indicator function which returns value 1 only if *m, n* ≥ 1). However, when the expressions *P*_*i(k*+1)_ and *P*_*α(k*+1)_ are zero or negative, this has the consequence that *no haplotype group* can fertilize these genotype groups. This means that the terms in the expected fitnesses *E*_*g*_ that contain terms *P*_*i(k+*1)_ and *P*_*α*(*k*+1)_ must be removed altogether. For *n* =1 female *S*_i(*k*+1)_ can never be fertilized, and for *n* = 0 no female with *S*_*k*+1_ can be fertilized. The same argument must be taken where incomplete haplotypes exist in the population. Furthermore, in general, the expected fitness in the population is 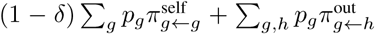 where we sum over groups g that can actually be fertilized, and is equal to 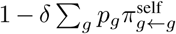 only if all groups can be fertilized.

## A.3 Results

We classify the results according to how many of the *k* resident haplotypes are complete (*n*). We start by recalling that whenever *m* = 0, i.e. *n* = *k*, then all *k* +1 haplotypes are complete, i.e. all haplotypes are maintained in the population and protected from extinction (protected coexistence).

### <u>**Case** *n* = 0</u>**: diversification is possible for** *k* = 3**, but never for** *k* > 3.

<u>*Proof:*</u> For *n* = 0, *m* ≥ 3 the above system simplifies considerably. As all fitnesses except *E*_*αβ*_ and *E*_α(*k*+1)_ are zero, and as the frequencies sum up to one, the dynamics are fully determined by a single difference equation, e.g. 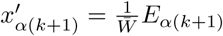, where

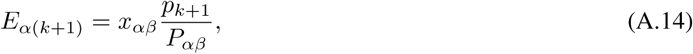

and where 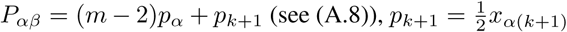 (see (A.13)) and the average fitness is 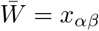 (because a female *S*_α_ *S*_β_ will be fertilized but *S*_α_*S*_*k*+1_ will not). The difference equation thus takes the form

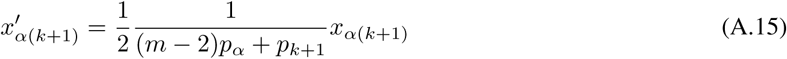

from which we can calculate the equilibria by setting 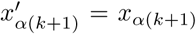. There exists only a single non-boundary (from now on non-boundary means that all frequencies are non-zero) interior equilibrium 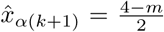 with as-sociated eigenvalue 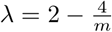. Clearly, the non-boundary equilibrium is an interior equilibrium (all frequencies are positive) only for *m* = 3, and for this value the eigenvalue is 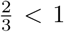, i.e. the equilibrium is locally asymptotically stable. We conclude that coexistence of three incomplete haplotypes and *S*_*k*+1_ is possible. For all *m* > 3 no interior equilibria exist and so coexistence is not possible. □

As shown above, interestingly, three incomplete SI haplotypes can coexist at a stable equilibrium together with one complete SI haplotype. This is surprising because female *S*_*k*+1_ can never be fertilized and so males would have to compensate for this fitness loss (see above). For *m* = 3, this actually is possible, because *S*_*k*+1_ -males can fertilize three out of six non-self genotype classes whereas resident *S*_*α*_-males can fertilize only one out of six non-self genotype classes. However, when m increases these proportions become more and more similar so that the advantage of *S*_*k*+1_ -males decreases enough for the haplotype *S*_*k*+1_ to go extinct.

### <u>**Case** *n* = 1</u>**: diversification is possible for** 3 ≤ *k* ≤ 6, **but never for** 7 **≤** *k*.

*Proof:* For *n* =1 genotype group *S*_*i*_*S*_*j*_ does not exist, and so we set *x*_*ij*_ = 0 and *E*_*ij*_ =0 in (A.1). Importantly, also *P*_*i*(*k+1*)_ =0 in (A.9), and so females *x*_*i*(*k*+1)_ remain unmated. The only complete haplotype that could fertilize *S_i_S_k_*_+1_ is part of the female genotype and is self-incompatible. We were not able to find an expression for an interior equilibrium for all *m* **≥** 2 and so we calculate the stability of boundary equilibria: whenever all boundary equilibria are unstable there is negative frequency dependent selection and all haplotypes will be able to coexist. The only (potential) boundary equilibria (see below) in this system are where 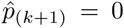 or 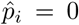 because only two SI haplotypes (*S*_*i*_, *S*_*k*+1_) can never persist in the system and so an equilibrium where *p*_*α*_ = 0 does not exist.

*The stability of* 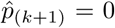: We calculate the dominant eigenvalue associated with an equilibrium where 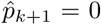, which in term of genotype group frequencies is 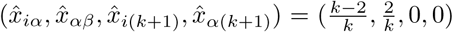. We take the Jacobian of the system, evaluate it at the resident equilibrium and obtain a dominant eigenvalue

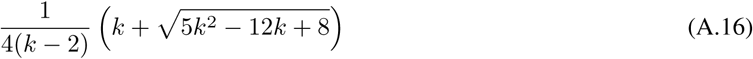

which is greater than 1 whenever 2 < *k* < 7.

*The boundary equilibrium* 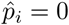: At this equilibrium haplotypes *S*_*α*_, *S*_*k*+1_ are the residents. From the previous case *n* = 0 we see that they can coexist for *m* = 3. Interestingly, however, the model with *n* =1 does not reduce to the model with *n* = 0 when we take the limit *p*_*i*_ → 0. This is because in *E*_*i*(*k*+1)_ (A.4) the term 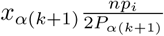, where *P*_*α*(*k*+1)_ = *np*_*i*_ doesn’t vanish for *p*_*i*_ → 0, and so it will always contribute in production of *x*_*i*(*k*+1)_, i.e the production of *S*_i_.

Since the only boundary equilibrium where *p*_*k*+1_ =0 is unstable whenever 2 < *k* < 7, this is when coexistence of the haplotype classes is possible. □

For *n* =1 the haplotype *S*_*k*+1_ can now be fertilized, but only when paired with *S*_*α*_ because there is only a single haplotype in group *S*_*i*_. In contrast to the previous case, females also contribute to fitness, but they are still at a selective disadvantage. For small *k* males can compensate for this, but similar to above, when *k* increases, the incomplete males *S*_*α*_ are able to fertilize an ever greater proportion of females. For sufficiently large *k* the haplotype *S*_*k*+1_ loses its advantage and the haplotype goes extinct.

### <u>**Case** 2 ≤ *n* ≤ *k*</u>: **long-term persistence of the diversification is not possible for any** *k* ≥ 3.

In this case, we can derive analytical results for any fixed *m* (we used *m* = 1,…, 10), but not for arbitrary *m*.

We found that for every *m* ≥ 2 the boundary where *p*_*k*+1_ = 0 has an associated dominant eigenvalue 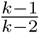, which is always greater than 1, and the boundary where *p*_*α*_ = 0 has associated dominant eigenvalue 1 (as in the previous case an equilibrium where only *p*_*i*_ = 0 is not a boundary equilibrium). However, we were unable to solve for all the equilibria as a function of *m* to exclude the possibility of stable interior equilibria. Nevertheless, we were able to solve the equilibria for specific values of *m* (we performed the calculations for *m* = 1,…, 10) and found that no interior equilibria exist (also, we have no reason why this should be any different for greater values of *m*). In addition, we performed numerical investigations to check that all trajectories approach the boundary equilibrium *p*_*α*_. The convergence is not asymptotical (dominant eigenvalue 1) and takes, approximately, 10^*x*^ generations for the frequency of *p*_*α*_ to be below 10^*-x*^ for any *x*.

In summary, we find that the equilibrium *p*_*α*_ = 0 has eigenvalue 1 while all other boundary equilibria are unstable and no interior equilibria exist. The convergence to the extinction of *S*_*α*_ thus takes a very long time, leaving the possibility for a diversification event if the incomplete haplotypes persist in the population long enough for (all of) them to gain the missing SLF *F*_*k*+1_.

<u>**Case** n = *k*</u>: All *k* + 1 SI haplotypes are complete and so negative frequency-dependent selection maintains the co-existence of all haplotypes.

**Necessary conditions for diversification.** If the above conditions are violated then no coexistence of *k* + 1 SI haplotypes is possible. Consequently, these conditions must necessarily be valid for a diversification to take place, but only when the (possible) SC intermediate haplotypes are excluded form the final composition of haplotypes (the condition for coexistence are only derived for SI haplotypes). Since intermediate SC haplotypes go extinct in pathways 2,3,4 we can apply our results there. In addition, it gives us a possible explanation for why pathway 1 is, in principle, possible for almost any combination of *n, m*. This is because the intermediate self-compatible haplotypes persist in the system throughout the diversification process and thus (apparently) decrease average fitness enough for the incomplete haplotypes not to be too disadvantaged that they go extinct. Also, this can be used for pathway 5 as this only has SI intermediates (see more discussion in Appendix C).

The above conditions, however, are not sufficient because (as discussed above) it is possible that even if *k* + 1 complete and incomplete haplotypes coexist, the necessary intermediate mutants, or the final mutant *S*_*k*+1_, are not able to invade the population. Interestingly though, the above conditions correctly predict whether diversification occurs in all cases except for a case where *n* = 0, *m* = 3 (relevant in pathway 4, see also below).

**Sexual selection: when are females selected against?** The above equations also reveal the similarities and differences between SR and NSR models in terms of sexual selection. In this paper, and similar to other models addressing SI systems, we have assumed that the population is well mixed (i.e. pollen disperses globally) and that each plant produces a large amount of pollen. Consequently, every female in the population will be fertilized if there exists at least one pollen grain that is able to fertilize it (i.e. compatible with that female). This is usually true in SR models where non-recognition results in compatibility, and so in SR models there is no sexual selection on females. Males, however, undergo frequency-dependent competition for fertilization and are under selection. In addition to sexual selection, individual fitness may be affected by inbreeding depression.

The situation is different in SLF based NSR models that involve incompleteness. A key feature in these models is that a male must be able to recognise a female, i.e. have the corresponding SLFs, in order to be able to fertilize this female. In such models, a novel female type might thus go unrecognised as no haploptype has yet the corresponding SLF. Thus, in contrast to SR models, in NSR models females can be under selection. This is the case, for example, in our model where haplotypes are assumed complete w.r.t all haplotypes except *S*_*k*+1_ (or *S*_*k*_ in cases where the mutants of *S*_*k*_ have not yet gained the SLF *F*_*k*_); if there are not enough complete SI haplotypes *S*_*i*_, females having *S*_*k*+1_ may not get fertilized. For *n* = 0 no females *S_x_S_k_*_+1_ (where *x* is any haplotype) can be fertilized, and for *n* =1 females *S_i_S_k_*_+1_ can never be fertilized because the only complete haplotype can’t fertilize itself. However, for n ≥ 2, all females can be fertilized (if we for now ignore the intermediate mutants who may lack *F*_*k*_). Therefore, only when n ≥ 2, there is no sexual selection acting on females, in which case all fitness effects come via males, or via inbreeding depression.

## B Diversification pathways 1 to 5: analytical and numerical results

To study whether diversification occurs for any of the pathways, we use (3) to construct the equations that correspond to each pathway separately. Then, for each pathway and possible initial state of the population we ask: can the first mutant invade the population? If so, will it coexist with all the haplotypes? Then, will the second (final) mutant invade and coexist with all the haplotypes? If these steps occur then this is diversification. There are in fact two possible final states; one where the intermediate (first) mutant is excluded, and one where it coexists with all initial and final SI haplotypes. It turns out that this distinction is important in SI models that allow for incompleteness because SC intermediate haplotypes influence the degree and nature of selection experienced by other, in particular incomplete, haplotypes (see Appendix C).

**Pathways 1 to 4.** Beyond the conditions derived in Appendix A (which we discuss in Appendix C) and the conditions derived for an analogous SR model of Uyenoyama *et al*. (2001) and Gervais *et al*. (2011), we did not find further analytical conditions concerning the feasibility of pathways 1 to 4; The conditions (analytical and numerical) for diversification pathways 1 and 2 are identical to pathways considered in Uyenoyama *et al*. (2001) and Gervais *et al*. (2011) when initially all haplotypes are assumed ’’complete” with respect to the not yet existing novel RNase (*n* = *k*). Moreover, for pathways 2,3 and 4 the first mutation is also identical to the pathways in Uyenoyama *et al*. (2001) and Gervais *et al*. (2011) for any *n* in our model, simply because the first mutation happens in the pollen and its fitness is therefore not affected by the presence or absence of the not yet utilized SLF. The second mutation however, which happens in the RNAse, does influence fitness differently for complete and incomplete haplotypes, consequently delineating our NSR model from the SR model in Uyenoyama *et al*. (2001) and Gervais *et al*. (2011). Similar to Uyenoyama *et al*. (2001) and Gervais *et al*. (2011), however, the outcome of the second invasion has to be computed numerically since we were not able to find an explicit solution for the intermediate equilibrium state. The complete numerical solution for their diversification events can be found in Figures S3-S5 (Supplemental Information) for three values of *k* = 3, 5,8 and all possible initial conditions 0 ≤ *n* ≤ *k*.

**Pathway 5.** The upcoming results, together with Appendix A, enable us to solve the pathway 5 (almost) fully analytically. The analysis is considerably simpler than for the other pathways because all haplotypes, including the intermediate haplotype, are all SI and so their fitness is not affected by inbreeding depression. To study diversification for pathway 5 we use (3) to analyze firstly whether the first mutant 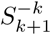 can invade the population, and if so, whether the second mutant *S*_*k*+1_ can invade and coexist with all the other haplotypes.

**Initial condition:** To consider pathway 5, at least one haplotype (the ancestral 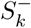) must be incomplete *m* ≥ 1.

**st Step:** In this paragraph we discuss the invasion of the incomplete mutant 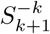 (i.e. mutant of the ancestral haplotype 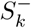that has the novel *R*_*k*+1_ but lacks *F*_*k*_ and *F*_*k*+1_) into a resident population for all *n, m*.

<u>*Case n* = 0</u>: (analytical result) Incomplete haplotype 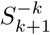is not able to invade the resident population for any *k*. This is simply because females of the mutant are never fertilized 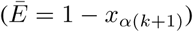, and the dominant eigenvalue is half, since males, when rare, have equal fitness compared to an average resident (a rare incomplete male with deficit one can fertilize as larger a proportion of females as can a common ”complete” resident, i.e. all but one haplotype class).

<u>*Case n* = 1</u>: (analytical result) Incomplete haplotype 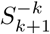 is not able to invade the resident population for any *k*. The reason is similar to above, except that now females can be fertilized, but only if paired with incomplete haplotypes *S*_*α*_, not when paired with complete haplotypes 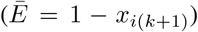. Females are thus at a disadvantage and since the contribution of males is as described in the previous case, the dominant eigenvalue is less than 1 (it is 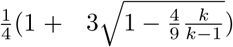 for all *k* ≥ 3).

<u>*Case* 2 < *n* < *k* - 1</u>: incomplete haplotype 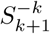 can ”neutrally” invade the resident population for any *k* ≥ 3 (dominant eigenvalue is one; analytical result) and then coexist with all the other haplotypes at a line of equilibria. We obtained the analytical expression for the line of equilibria for the cases *n* = 2, *m* =1,2 and *n* > 2, *m* =1, and for the remaining cases *n* > 1, *m* ≥ 2 we found it numerically.

The dominant eigenvalue is 1 because now mutant females can be fertilized when paired with both *S*_*i*_ and *S*_*α*_ 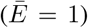, and males are as described in the previous two cases. The line of equilibria is a consequence of the fact that the mutant 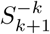.and its ancestor 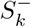 have exactly the same fertilization properties: all females can be fertilized, and males can fertilize exactly the same haplotype classes, all but their own and each other. They are thus selectively neutral w.r.t each other, and the only force that can change their relative frequencies is drift in finite populations.

**2nd Step:** Here we assume that the first mutant 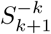 and its ancestor 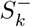 can coexist long enough for the second

(final) mutant *S*_*k*+1_ to appear in the population (case *n* ≥ 2). In principle we should evaluate the invasion ability at the line of equilibria, which can in some cases be calculated (see above), but we take the simpler route and assume that the first mutant is still rare (negligible frequency) by the time *S*_*k*+1_ arrives in the population. In such a scenario we can apply the results from section A. Following this, for all *n* ≥ 2, *k* ≥ 3 the novel mutant *S*_*k*+1_ can invade the population (we also expect this result to hold when evaluating the population at the line of equilibria). However, as discovered in A, for *n* ≥ 2, *k* ≥ 3 all incomplete haplotypes are at disadvantage and eventually go extinct, but, since the extinction is very slow the corresponding haplotype classes can be rescued by completing the haplotypes (see above).

## C Discussion on the necessary conditions A and the analytical and numerical results on pathways 1 to 5

Firstly, even though it seems like pathway 1 provides the most compelling parameter region (α, δ, *n, k*) for diversification, for the very same parameter region the other pathways 2, 3, 4 lead to full extinction of all SI haplotypes, and replacement by a single SC haplotype. This suggests, that, if the population experiences favourable conditions for diversification for pathway 1, any mutation that leads to a complete SC haplotype (pathways 2,3,4) will result in the loss of incompatibility from the population. Secondly, we may argue that the diagonal mutations where a specific SLF *F*_*k*+1_ mutates to another specific SLF *F*_*k*_ is approximately *L* times less likely than any other mutation on the cube, thus taking a longer time to occur. We thus predict that pathway 1 (but also pathway 2) is unlikely to be responsible for diversification.

We have shown that diversification via pathways 2, 3 and 4 is possible for *n* = *k* and for *n* = 1 and 3 ≤ *k* < 7, in which case (long-term) stable coexistence occurs for *k* + 1 SI haplotypes, *k* - 1 of them incomplete (missing *F*_*k*+1_). Interestingly, the coexistence of complete and incomplete haplotypes is long-term, until one of the incomplete haplotypes becomes complete, resulting in altogether three complete SI haplotypes, which then drive all the remaining incomplete haplotypes to extinction. However, this happens very slowly and so the incomplete haplotypes (”classes”) can be rescued by gaining the missing SLF *F*_*k*+1_ before the class goes extinct. Nevertheless, unless all incomplete haplotypes become complete, the diversification process will result in the destruction of incomplete haplotypes and the number of surviving haplotypes drops to the number of complete haplotypes in the current population (which is likely to be lower than the number in the initial resident population *k*). This diversification path may therefore eventually lead to a reduction in haplotype classes.

If the initial resident population (pathways 2 to 4) were to have at least two complete haplotypes *n* ≥ 2, then immediately after the invasion of *S*_*k*+1_ all incomplete haplotypes proceed towards extinction (and we are back to the above situation). The diversification is thus only short-term, and will not persist unless, as above, all incomplete SI haplotype classes are rescued by gaining the missing SLF *F*_*k*+1_. We thus predict, that for small *k* diversification happens usually for *n* =1 since long-term stable coexistence is possible; while for higher k and/or greater *n* diversification may occur, but only for higher mutation rates. In addition, we should observe sudden drops of the number of haplotype classes associated with the creation of new complete SI classes.

Pathway 5, interestingly, is free from inbreeding depression during the diversification process because all individuals outcross. However, our analytical conditions predict that if diversification happens, it is very likely only short-term, unless the mutation rate is sufficiently high for the incomplete haplotypes to become complete. Moreover, at least two haplotype classes need to have the not yet utilized SLF to initiate the diversification process.

